# Swine Growth Promotion with Antibiotics or Alternatives can Increase Antibiotic Resistance Gene Mobility Potential

**DOI:** 10.1101/2020.02.19.957100

**Authors:** Johanna Muurinen, Jacob Richert, Carmen Wickware, Brian Richert, Timothy A. Johnson

## Abstract

Even though the use of antibiotics for food-producing animals may contribute to the emergence of antimicrobial resistance, antibiotics are still used as growth promoters. Due to consumer and regulatory pressures, the use of alternatives to antibiotics as growth promoters is increasing, thus more information is needed on their capability to disseminate antimicrobial resistance compared to antibiotics. We investigated the impacts of carbadox (antibiotic), copper sulfate and zinc oxide (metals) and mushroom powder (natural product) on the pig fecal resistome and microbiome. Antibiotic resistance gene (ARG) and mobile genetic element (MGE) abundances were measured using a high-throughput qPCR array with 382 primer pairs. Bacterial community composition was determined by 16S rRNA gene sequencing. More ARGs co-occurred with MGEs in the growth promoter group samples than in the control group samples. Community composition could not be linked to resistome in the growth promoter group samples, indicating a potential decoupling of ARGs and phylogeny. Additionally, machine-learning methods aided in defining the community and resistome differences in response to treatments. Since increased ARG mobility potential was the primary response to the dietary additives used in this study, we suggest that ARG mobility should be considered when designing antimicrobial use policies and antimicrobial resistance surveillances.

## Introduction

Antibiotics have been used in pork production to prevent diseases and to increase productivity since the 1940’s^1^. Due to the established connection of antibiotic use in animals and the emergence of antibiotic resistance in pathogenic bacteria^2^, the regulation and prohibition of antibiotic growth promoters is increasing (prohibition in the EU in 2006^1^, restrictions in 2017 in the U.S.^3^, and restrictions in 2020 in China^4^). Additionally, a report to the Secretary General of the United Nations suggested that antibiotic growth promoters should be completely phased out from livestock production^5^. Despite the increasing regulation, most of the world’s pork is produced in countries that still allow growth promotion with antibiotics. In 2020, the U.S. was the third largest producer of pork worldwide after China and the EU, and the second largest exporter of pork meat after the EU. Currently, the livestock industries in the U.S. as well as in many other countries are looking for alternative feed additives to replace antibiotics as growth promoters due to increasing restrictions and consumer concerns towards antibiotic use in animal agriculture. Because of these reasons, the markets are now favorable to so-called alternatives to antibiotics as growth promoters and the interest towards them is increasing rapidly^6,7^.

Among the antibiotics that are still allowed for growth promotion in the U.S., carbadox is used in pigs mainly to control dysentery and bacterial enteritis. Carbadox can cause short-term but also long-lasting alterations in the microbiome^8^ and can promote the mobility of antibiotic resistance genes (ARGs) through transduction^9^. Still, little is known about the impact of carbadox or especially alternative feed additives on the abundance of ARGs in pig feces. Numerous growth promoters that are alternatives to antibiotics are currently under study for their efficacy to promote animal growth and gastrointestinal health. Among natural products, of special interest are medicinal mushrooms. *Cordyceps militaris* and *Ophiocordyceps sinensis* (formerly *C. sinensis*) produce many bioactive compounds, such as Cordycepin and Beta-Glucan, which have antimicrobial effects and can enhance the immune system^10^. Zinc (Zn) and copper (Cu) are important trace elements for all organisms, and livestock animals commonly receive feed supplements that ensure their required concentration in the feed. Higher concentrations of Zn and Cu can be used for controlling various bacterial infections in livestock animals, such as diarrheal diseases^7^ and also for growth promotion^7,11,12^. However, use of Zn oxide has shown to select zoonotic methicillin resistant *Staphylococcus aureus* (MRSA) and multiresistant *E. coli*^13–16^. Consumption of Cu-supplemented animal feed can increase the prevalence of erythromycin resistance in gram-positive bacteria^17^ and various plasmid-mediated resistance genes in microbiomes^16,18^. Thus, it seems that some growth promoters that are alternatives to antibiotics are actually antimicrobials and may select ARGs similarly as the antibiotics they are meant to replace.

On the other hand, when quantifying the antibiotic resistome, it is important to consider that changes in resistance gene abundance may simply be due to a change in the underlying community composition, since community composition can explain resistance gene composition^19,20^. It is thought that growth promoters alter the gut microbial community composition, and that the microbiota composition could be manipulated to promote animal growth by inducing populations favorable to growth, immunity, and gut health^21^. Therefore, it is important to characterize the bacterial community composition to determine if the populations linked to induced animal growth carry ARGs and mobile genetic elements (MGEs). However, making connections between taxonomic data and resistance gene data can be challenging, since microbiome datasets are compositional and rarely meet assumptions of normality that many statistical tests require^22,23^. In addition, it has been pointed out that different normalization strategies could influence the results derived from sequence data^24,25^.

In order to examine the influence of carbadox, cordyceps mushroom powder and pharmacological concentrations of Zn and Cu on the community composition and resistome^26^, we took fecal samples from pigs that were administered these growth promoters, extracted DNA, analyzed 16S rRNA gene sequences and quantified genes related to resistance and horizontal gene transfer with 382 primer pairs using a high-throughput qPCR array^27,28^. We also compared two normalization strategies for taxonomic data using statistical analyses that are suitable for compositional datasets. Our results suggest that inclusion of ARGs into MGEs led to decoupling of bacterial community composition and resistome composition in response to growth promotion, highlighting the importance of MGEs in shaping the resistomes under the influence of different types of antimicrobial agents.

## Results

### Samples and data quality control

Samples were collected from pigs that had been assigned into non-treatment control group (NTC = no antibiotic or alternative growth promoters), or one of the growth promoter groups (AB = carbadox, M = mushroom powder mixture of *C. militaris* and *O. sinesis,* ZnCu = Zn oxide and Cu sulfate). Each treatment group consisted of six pens and each pen had seven pigs. The DNA for 16S rRNA gene sequencing was extracted from combined fecal samples of one medium weight female and male per pen. After quality filtering, a total of 741,785 sequences were obtained. For 22 samples sequences per sample ranged from 11,392 to 67,072. Two samples were discarded due to low number of sequences, resulting in five samples in the NTC and AB groups. The data analysis for 16S rRNA gene sequencing was completed using two different methods: rarefication and subsampling or total sum scaling (TSS). The TSS normalized 16S rRNA gene sequence data had 132 operational taxonomical units (OTUs), while rarefied and subsampled 16S rRNA gene sequence data 127 OTUs. The same DNA samples were used for qPCR array analysis to examine how the treatments altered the resistome. One hundred and thirty-six assays out of 382 assays (Supplementary Table S1) targeting antibiotic resistance genes (ARGs) or mobile genetic elements (MGEs) were positive. See materials and methods for qPCR data processing and Ct value adjustment for four assays that had unspecific amplification.

### The profiles of most abundant genera and genes were similar in different treatment groups

To take into account the influence of normalization of 16S rRNA gene sequence reads on the results, we used two different normalization methods. Only small differences were discovered between TSS normalized and rarefied and subsampled OTUs among the most abundant genera (Fig. 1A and B). The TSS normalized OTUs and rarefied and subsampled OTUs correlated significantly (ρ = 0.97, *p* < 0.05) (Supplementary Fig. S1), indicating the agreement of the overall community composition of the two normalization methods. *Prevotella* was the most abundant genera in all treatments and all samples had several short chain fatty acid producers (Fig. 1A and B). Very few OTUs were found only in one treatment group with both normalization methods and a majority of the OTUs were found in all treatment groups, however more OTUs were found in all treatment groups using TSS normalization than rarefied and subsampled normalization (Fig. 1D and E).

**Fig. 1.**
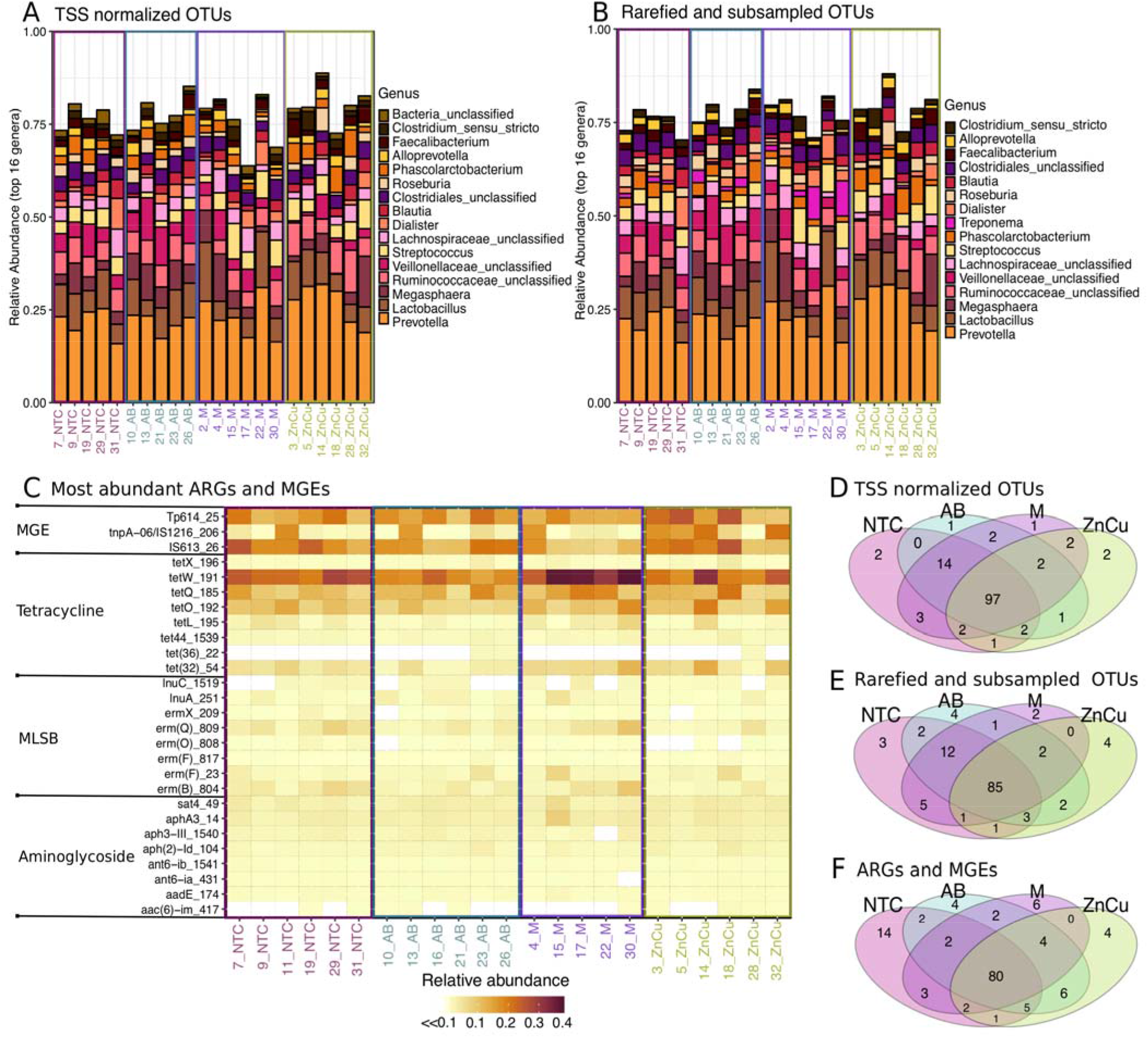
Comparison of most abundant genera, ARGs and MGEs in different treatment groups. Samples on x-axis are grouped according to the treatments and color-coded. Sample names are as follows: NTC= Non-treatment control, AB= Carbadox (antibiotic), M= Mushroom powder and ZnCu= zinc oxide and copper sulfate. The number in front of the group code denotes the number of the pen. (A) Stacked bar plot showing 16 most abundant genera in OTUs normalized using total sum scaling (TSS). (B) Stacked bar plot showing 16 most abundant genera in rarefied and subsampled OTUs. (C) Most abundant genes related to antibiotic resistance and mobile genetic elements (n=27). Each row represents results of each primer set (assay) (Supplementary Table S1) displayed on the y-axis. Assays are grouped according to the antibiotic group which the target genes confer resistance. MLSB is abbreviation for Macrolide-Lincosamide-Streptogramin B resistance and MGE for mobile genetic elements. One sample from mushroom powder group was left out from the qPCR array analysis due to a technical error. (D) Venn diagram showing the OTUs that are shared between samples belonging into different treatment groups when TSS normalization was used. (E) Venn diagram showing the OTUs that are shared between samples belonging into different treatment groups when Rarefying and subsampling was used. (F) Venn diagram showing the ARGs and MGEs that are shared between samples belonging into different treatment groups.

The different treatment groups also had similar resistome profiles (Fig. 1C). The NTC group had the highest number of positive assays (110), the AB group had 106 positive assays, The M group had 100 positive assays and the ZnCu group had 103 positive assays. Twenty-eight genes were detected in only one treatment group, but most genes were found in all treatment groups (Fig. 1F). Out of the positive assays, 108 targeted ARGs and 28 MGEs. Among the detected ARGs, 63 conferred resistance to aminoglycosides, MLSBs or tetracyclines and the most common resistance mechanism was antibiotic deactivation (Fig. Supplementary S2A and B). Among MGEs, most of the positive assays targeted insertion sequences (12) or transposases (7) (Fig. Supplementary S2C).

### Growth promoters favored different genera, ARGs and MGEs

Generalized linear models (GLMs) with False discovery rate control^29^ for *p*-value adjustment were used for testing which genera, ARGs and MGEs differed in abundance between treatments. Mostly only minor differences in abundance of different genera were found between treatment groups (Supplementary Fig. S3A and B). Compared to all other growth promoters, carbadox favored unclassified *Veillonellaceae* in feces and decreased the abundance of *Streptococcus,*whereas Zn and Cu decreased the abundance of *Bifidobacterium* and *Campylobacter* (adjusted *p*-value□<□0.05, gamma distribution GLMs [TSS OTUs] and negative binomial GLMs [rarefied and subsampled OTUs]) (Fig. 2A and B, Supplementary Table S2 and S3). Mushroom powder decreased the abundance of fecal *Roseburia* and favored *Campylobacter* compared to other treatments (adjusted *p*-value □<□0.05, gamma distribution GLMs [TSS OTUs] and negative binomial GLMs [rarefied and subsampled OTUs]) (Fig. 2A and B, Supplementary Table S2 and Supplementary S3). *Streptococcus* was more abundant in M and ZnCu group samples compared to other groups (adjusted *p*-value □<□0.05, gamma distribution GLMs [TSS OTUs] and negative binomial GLMs [rarefied and subsampled OTUs]) (Fig. 2A and B, Supplementary Table S2 and S3). More ARGs and MGEs were differentially abundant in NTC group and M group comparison than when NTC group was compared to ZnCu or AB groups (Supplementary Fig. S3C). Compared to all other growth promoters, carbadox and Zn and Cu increased the relative abundance of *vat(E),* mushroom powder increased the relative abundance of *tetW,* and Zn and Cu favored *tetM* (adjusted *p*-value □<□0.05, gamma distribution GLMs) (Fig. 2C, Supplementary Table S4). Interestingly, carbadox treatment decreased the relative abundance of *tetW, tet(32), erm(B), ermT* and *tetM* compared to other treatments or to NTC group (adjusted *p*-value □<□0.05, gamma distribution GLMs) (Fig. 2C, Supplementary Table S4).

**Fig. 2.**
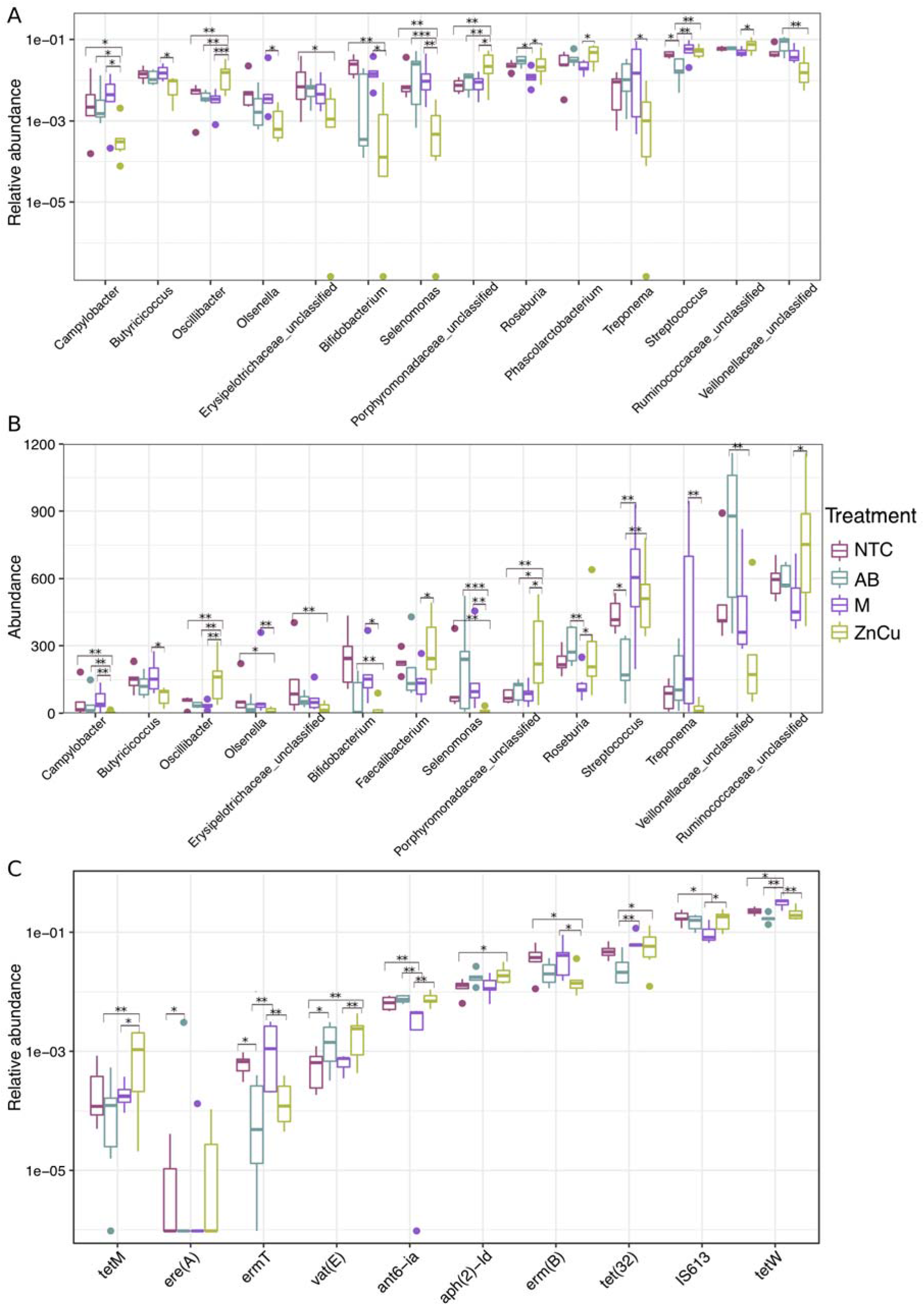
Boxplots showing the most abundant genera, ARGs and MGEs with statistically significant differences between treatment groups labeled on right. NTC= Non-treatment control, AB= Carbadox (antibiotic), M= Mushroom powder and ZnCu= zinc oxide and copper sulfate. The asterisks “*”, “**” and “***” denote statistical significance levels at *p* < 0.05, *p* < 0.01 and *p* < 0.001, respectively. (A) Most abundant genera in TSS normalized OTUs (n=14), (B) Most abundant genera in rarefied and subsampled OTUs (n=14), (C) Most abundant genes related to resistance and transfer (n=10). See Supplementary Fig. S3A, B and C for all differentially abundant genera and ARGs and MGEs and Supplementary Table S2, S3 and S4 for fold changes of the differently abundant genera and ARGs and MGEs.

Both OTU normalization methods yielded similar results for differential OTU abundance in the treatment groups (Fig. 2A and B). The exceptions were that *Phascolarctobacterium* was among the differentially abundant genera when TSS normalization was used, whereas *Faecalibacterium* was differentially abundant when rarefying and subsampling was used (Fig. 2A and B) and that more genera were differentially abundant when TSS normalization was used (Supplementary Table S2 and S3).

### Growth promoters had modest influences on overall bacterial community composition and resistome

With both TSS normalized OTUs and rarefied and subsampled OTUs, the treatment group explained 24% of the variability in community composition (PERMANOVA, R^2^= 0.239, *p* < 0.05). With ARGs and MGEs, the treatment group explained 27% of the variability (PERMANOVA, R^2^ = 0.267, *p* < 0.05). The number of sequences per sample was also included as a variable in the PERMANOVA model with both normalization methods, but the library size did not have influence on the community variability in our samples (*p*> 0.05).

The different growth promoters altered the community composition and resistome only slightly, since in all non-metric multidimensional scaling (NMDS) ordinations the samples belonging to different treatment groups clustered close to each other with mostly overlapping centroid confidence intervals (Fig. 3A, B and D). In ordinations of TSS normalized and rarefied and subsampled OTU-tables, the ZnCu group tended to cluster further away from the other samples indicating more dissimilar community composition (Fig. 3A and B), however there were no significant differences between treatment groups among pairwise PERMANOVA comparisons (False discovery rate control adjusted^29^ *p*-value > 0.05). In the ordination of the ARG and MGE data, there was no separation between ZnCu group and the other groups, instead the M group clustered slightly separately (Fig. 2D) and in the pairwise comparisons using the ARG and MGE data, M vs. AB group explained 32% of the variance (PERMANOVA, R^2^ = 0.321, False discovery rate control adjusted^29^ *p*-value < 0.05), but in all the other pairwise comparisons the differences were nonsignificant (False discovery rate control adjusted^29^ *p*-value > 0.05).

**Fig. 3.**
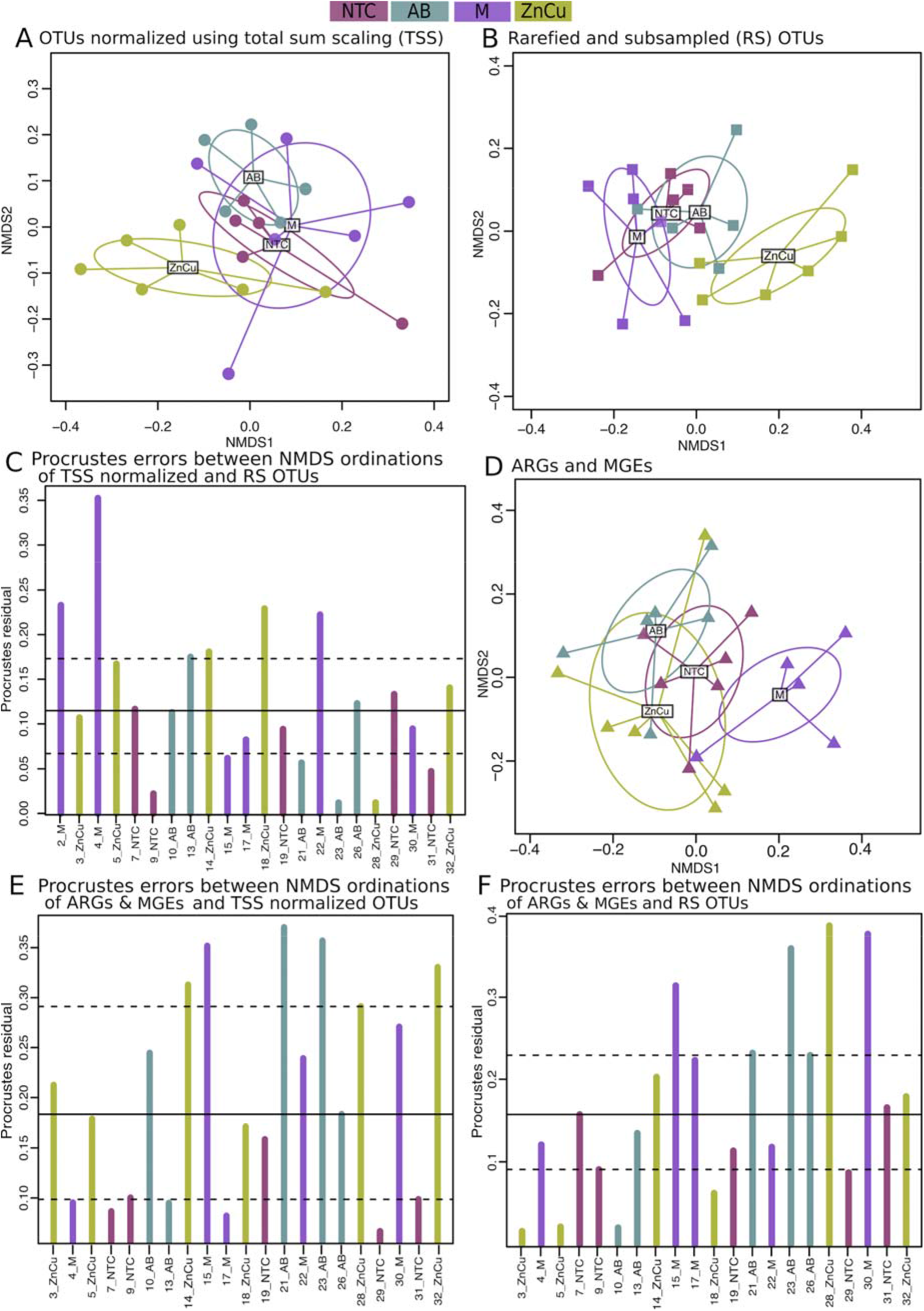
NMDS ordinations of TSS normalized OTUs, rarefied and subsampled OTUs and ARGs & MGEs and Procrustes errors between them. Sample names are as follows: NTC= Non-treatment control, AB= Carbadox (antibiotic), M= Mushroom powder and ZnCu= zinc oxide and copper sulfate. The number in the sample name denotes the number of the pen. (A) NMDS ordination of TSS normalized OTUs. (B) NMDS ordination of rarefied and subsampled (RS) OTUs. (C) Procrustes errors between NMDS ordinations of TSS normalized OTUs and rarefied and subsampled OTUs. (D) NMDS ordination of ARGs and MGEs. (E) Procrustes errors between NMDS ordinations of TSS normalized OTUs and ARGs & MGEs. (F) Procrustes errors between NMDS ordinations of rarefied and subsampled OTUs and ARGs & MGEs. The Procrustes residual error line plots (C, E and F) allow residual error size comparisons. The bars show the difference in the community structures between the two normalization methods (C) as well as the differences in community structure and resistome structure in samples belonging to different treatment groups (E and F). Horizontal lines denote the median (solid), 25% and 75% quantiles (dashed).

### Taxonomic variation did not explain resistome variation in growth promoter group samples

The OTU ordinations were correlated against each other and against the ARG and MGE NMDS ordination using Procrustes analysis to determine if the taxonomic variation explains resistome variation. The two OTU NMDS ordinations had reasonably high (0.7) and significant (*p* < 0.05) correlation and the Procrustes residual error remained lower than 0.25 in all samples except one (Fig. 3C). The correlation between the rarefied and subsampled OTU ordination (which did not include all the data due to subsampling) and the ARG and MGE ordination was moderately high (0.6, *p* < 0.05) and there was no pattern in Procrustes residual errors across different samples (Fig. 3F). Contrariwise, the correlation between the TSS normalized OTU ordination (that included all quality filtered data) and the ARG and MGE ordination was nonsignificant (*p*> 0.05) and Procrustes residual errors were high in most samples not belonging to NTC group, in which the residual errors were mostly equal or less than the first quantile residual value (Fig. 3E). The Procrustes analyses performed on TSS normalized OTUs and resistome implies that taxonomic variation explained resistance variation in NTC samples with less error. However, growth promoters altered the resistome, resulting higher residual errors in samples belonging to AB, M and ZnCu groups and thus the overall correlation between TSS normalized OTU ordination and ARG and MGE ordination was not significant (Procrustes analysis, *p* > 0.05).

The links between bacterial community structure and resistome were further examined with Mantel’s test using Spearman’s rank correlation. The correlation coefficients between the TSS normalized OTUs and ARGs and MGEs distance matrices as well as the rarefied and subsampled OTUs and ARGs and MGEs distance matrices were low (ρ = 0.25 and ρ = 0.23, respectively, *p* < 0.05), which suggests that the phylogenetic composition did not govern the resistome composition when all samples were included in the analysis. We also used Mantel’s test for all treatment groups individually. With the NTC group and both OTU normalization methods, the bacterial community distance matrix correlated significantly with the distance matrix obtained from ARGs and MGEs (ρ = 0.66 [TSS OTUs] and ρ = 0.62 [rarefied and subsampled OTUs], *p* < 0.05). With all growth promoter groups and both OTU normalization methods, the correlations between OTU and ARG and MGE distance matrixes were nonsignificant, giving more evidence that the growth promoters shaped the resistance composition and thus taxonomic variation did not explain the resistance variation in the growth promoter group samples. It should be denoted that we had only five or six samples in each treatment group; however, the results were consistent, since the correlations between taxonomic structure and resistome were reasonably high and significant in NTC group and nonsignificant in all other groups.

### Growth promoters increased the co-occurrences of ARGs and MGEs

A correlation matrix between ARG and MGE relative abundances was visualized using network analysis to examine if the growth promoters selected resistance genes into mobile genetic elements. The network of NTC group was simpler compared to growth promoter group networks as the number of correlating ARGs and MGEs increased in response to growth promotion (Fig. 4). There were no integrons co-occurring with resistance genes in the NTC group network, but integrons were present in all the growth promoter group networks (Fig. 4) and co-occurred with many ARGs. AB and M group networks had more aminoglycoside resistance genes than other networks and M group also had the most multidrug resistance genes, but the least tetracycline resistance genes, whereas AB group network had the most vancomycin resistance genes (Fig. 4).

**Fig. 4.**
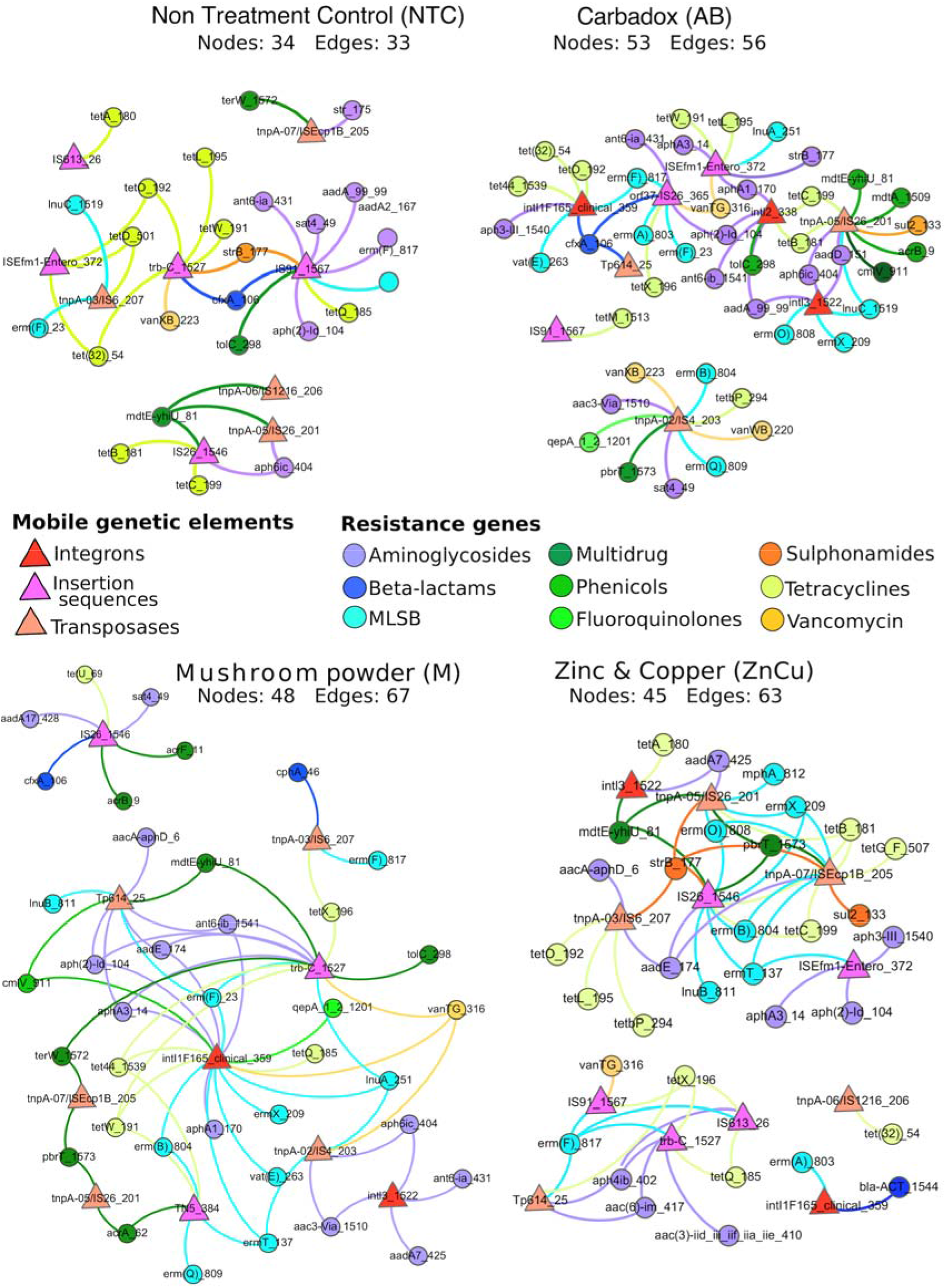
Network analysis showing co-occurrence patterns between ARGs and MGEs within the samples in different treatment groups. Nodes of the MGEs are triangles, and circle resistance gene nodes are colored according to the antibiotic they confer resistance. Edges between resistance gene nodes and mobile genetic element nodes have the color of the resistance gene node. Nodes have equal sizes, edges have equal weights, and distance between the nodes is irrelevant.

### Transposase gene for insertion sequence-like element IS1216 was the best predictor for resistance and community composition

We used machine-learning approaches to identify drivers of changes in community composition. First, a cluster analysis was performed on a combined data table of TSS normalized OTUs and ARGs and MGEs using *t-*distributed stochastic neighbor embedding (t-SNE) algorithm and HDBSCAN algorithm^30^. Then, a classification random forests model ^32,33^ was used for identifying the predictors for the clustering pattern. Three clusters were identified after dimension reduction. Most of the samples belonged to cluster 2, while cluster 1 had two ZnCu samples, one M group sample and one AB group sample, and cluster 3 contained only three ZnCu samples (Fig. 5A). According to the partial dependence plot that shows the most important predictors found by the classification random forests model, clusters 1 and 3 were separated from cluster 2 due to higher abundance of transposase gene linked to IS1216 element and from each other because the abundance of *tetO* was lower in cluster 3 (Fig. 5B). Five of the nine best predictors for clustering pattern were MGEs and ARGs and four were bacterial genera (Fig. 5B). Interestingly, only one of the ZnCu group samples belonged to cluster 2, while all the other ZnCu samples clustered to clusters 1 and 3. Thus, growth promotion with Zn and Cu might cause more alterations in the community composition and resistome than the other growth promoters examined in this study.

**Fig. 5.**
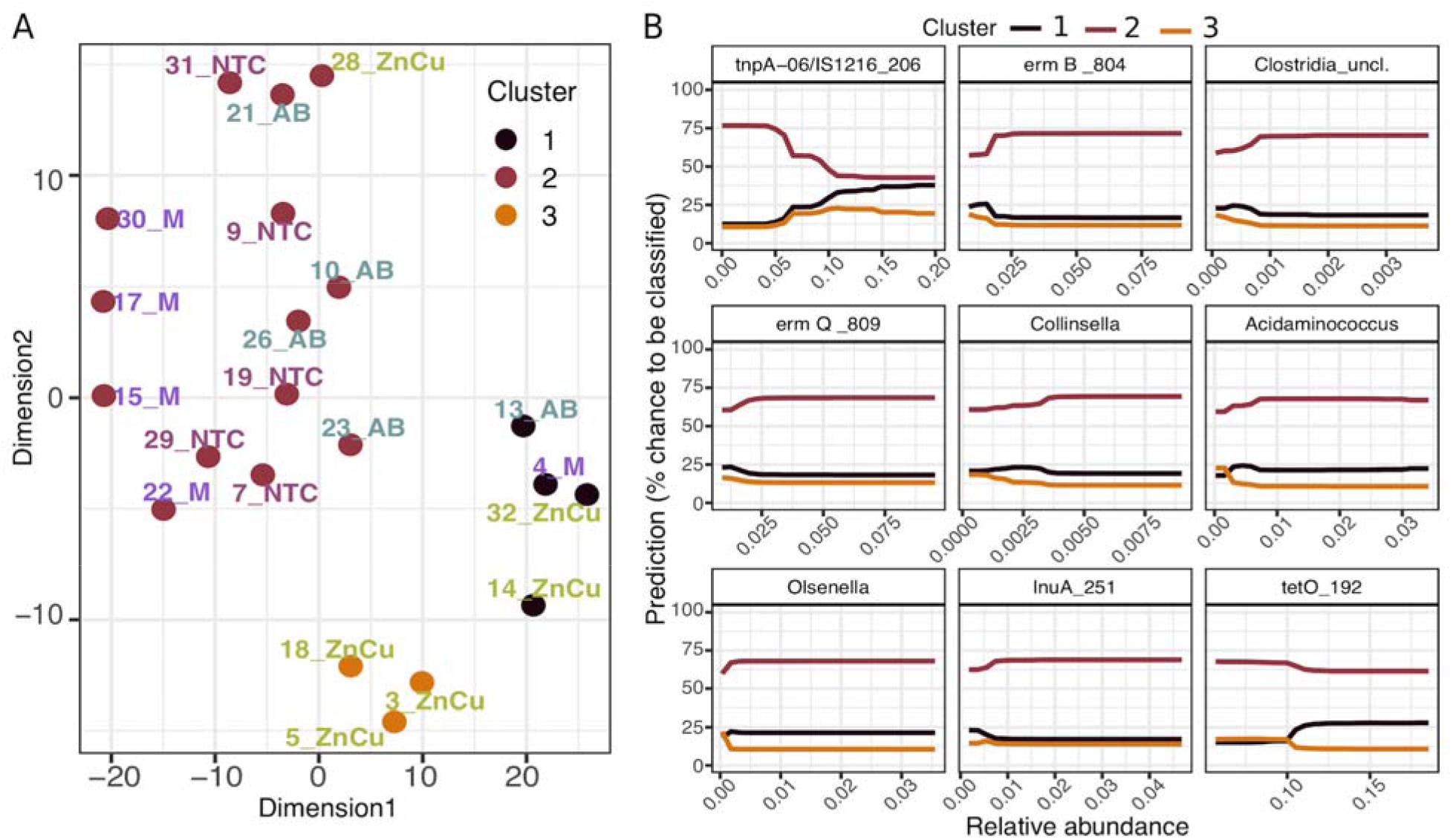
t-SNE analysis of a dataset containing ARGs, MGEs and TSS normalized OTUs (all relative abundances) and the distribution of the pens where experimental pigs were kept. (A) The clustering pattern of the samples. (B) Partial dependence plot of cluster numbers and the most important predictors in the order of decreasing importance. The partial dependence plot shows the effect of each predictor on the model outcome one by one, while the other predictors are fixed to their average value.

## Discussion

Our study explored the influences of antibiotic and alternative growth promoters on the pig fecal resistome, ARG mobility, and bacterial community composition. Resistome structure and taxonomic structure correlated in the NTC group samples but there was no correlation between resistome and phylogeny in the growth promotion groups. Some have reported that community composition predicts resistance profile^19,20,34^, while others have reported that under the presence of a selective pressure, phylogeny and resistome can become uncoupled in the swine feces and manure^35^. Our network analysis revealed that the growth promoters in this study increased linkages between ARGs and MGEs, i.e. more ARGs correlated with MGEs in the resistome of growth promoter group pigs than in the resistome of NTC group pigs. Thus, it seems plausible that resistance gene mobility would have increased in response to growth promotion, since subinhibitory antimicrobial concentrations and bacterial stress response mechanisms can induce horizontal gene transfer^36,37^.

Besides the inclusion of ARGs into MGEs, another possible scenario is that bacteria carrying MGEs with multiple ARGs and other embedded features would have tolerated the growth promoters better than bacteria carrying MGEs with only few ARGs, possibly due to more variability in stress response mechanisms^16,36,37^. Despite the mechanism, more ARGs co-occurring with MGEs in response to growth promotion could indicate more persistent resistance gene collection^38^ and potential for ARGs to be mobilized by the MGEs in the future, and this way the ARGs in the animal microbiomes could eventually be transmitted to human microbiomes^2,36^.

Despite the changes in ARG mobility potential, the abundances of nearly all ARGs and MGEs were on similar levels in the non-treatment control group and the growth promoter groups, which indicates that it is unlikely that changing the substance that is used for growth promotion would reduce antimicrobial resistance in the pig fecal microbiome, at least in a short period of time (33 days in this study). Gut bacteria, especially gram-negative species, are known to carry many ARGs and MGEs and the gastro-intestinal tract is suspected to be a major hotspot for horizontal gene transfer^36,39^. These observations have also been made with individuals without antibiotic exposure^40,41^.

Interestingly, some ARGs were less abundant in the AB group than in the NTC group and alternative growth promoter groups. A potential explanation for this could be that carbadox is a broad acting antibiotic and suppresses most bacterial populations^8^. Overall, we did not observe large shifts among genera as a result of growth promotion feed additives. *Bifidobacteria,* which have been previously linked to lower antibiotic resistance level^42,43^, were somewhat more abundant in the NTC and M group samples; however, in this study the ARG abundances in samples belonging to different groups were similar. It is possible that the shifts in the community composition caused by growth promoters were small with high variability and thus the changes in the community structure are difficult to capture with community-wide molecular approaches.

Although the changes in community composition and resistome in response to treatments were modest and mostly nonsignificant, we observed shifts with machine learning methods. The differences we observed did not precisely follow the experimental grouping (but largely isolated the ZnCu group), and therefore they were not found with commonly used ordination methods or differential abundance analysis. The transposase gene linked to IS1216 element was the driver for the clustering result and has been previously associated to Gram-positive bacteria^35,44^, mainly *Enterococcus faecium*^45^. However, *Enterococcus* was not among the most abundant or differentially abundant genera, or among the predictors of the clustering result in random forest analysis. Even though we cannot demonstrate all the factors explaining the clustering result, the observation that zinc and copper treatment perhaps altered the microbiome and resistome more than the other treatments would not have been possible without the use of a machine learning approach for statistical analyses. Thus, machine learning is an important tool in determining the complex relationship between resistome and bacterial community compositions.

Considering the two approaches to normalize OTU counts between samples, the rarefied and subsampled OTU data and TSS normalized OTU data mostly agreed; however, in Procrustes analysis, the rarefied and subsampled OTU data correlated significantly with the resistome data when all samples were used, although the same analysis using the TSS normalized data as input showed that the taxonomic data did not explain resistance. It is important to acknowledge that rarefying and subsampling captures the most abundant OTUs for each sample^25^ and discards rare OTUs, as well as slightly adjusting OTU relative abundances in different samples. If the sample size and differences in OTU abundances are small before the procedure, the shifts might change the outcome if rarefied OTU data is used in comparison with data obtained using a different method. Thus, researchers making connections between taxonomic data and resistome observations should use multiple methods to confirm their findings.

In this experiment, we observed a decoupling of phylogeny and resistance gene composition, possibly through increased co-occurrences between ARGs and MGEs. This indicates that the studied feed-additives can increase and maintain resistance gene mobility potential. On the other hand, under this experimental design, the withdrawal of antibiotics did not decrease the abundance of antibiotic resistance, however, we did not observe enrichment of ARGs in response to growth promotion either. Thus, we suggest that the mobility potential of ARGs should be considered to include in antimicrobial resistance surveillances to establish the impact of changes in agricultural practices on antimicrobial resistance.

## Materials and methods

### Animal experiment statement

All procedures involving animal use were approved by the Purdue University Animal Care and Use Committee (protocol #1303000841), and animal care and use standards were based upon the Guide for the Care and Use of Agricultural Animals in Research and Teaching^46^.

### Samples and DNA

The pig fecal samples were obtained from growth promoter experiment where 210 weanling pigs ((Duroc × (York × Landrace)) avg. 19 d of age and 5.8 kg were used in a 33-day trial. The experiment had 7 pigs in each pen and 6 pens per each treatment. Feed amendment treatments were: 1) non-treatment control (NTC); 2) antibiotic growth promoter (carbadox, 55 ppm) (AB); 3) mushroom powder (mixture of *C. militaris* and *O. sinesis,* 300 ppm) (M); 4) carbadox and mushroom powder mixture (results are not included in this study); 5) copper sulfate (125 ppm) and zinc oxide (3000 ppm d 0-7, 2000 ppm d 7-35) (ZnCu). After 33 days, fecal samples were taken from 1 median weight female and male per pen. Samples from the same pen were pooled, and DNA was extracted using the DNeasyPowerLyzer PowerSoil DNA Isolation Kit (Qiagen) according to the manufacturer’s protocol. Extracted DNA was stored at −20 °C before 16S sequencing and qPCR array.

### 16S sequencing and quantitative PCR array

The 16S rRNA gene library was constructed as described^47^. Briefly, the V4 region of the bacterial 16S rRNA gene was amplified with the 515R (GTGCCAGCMGCCGCGGTAA) / 806R (GGACTACHVGGGTWTCTAAT) primers. 16S rRNA gene libraries were also prepared for a known mock community (20 Strain Even Mix Genomic Material; ATCC® MSA–1002TM) and a no-template control (water). The amplified DNA from one 96-well plate was normalized using a SequalPrep Normalization Plate (Invitrogen), and pooled into a single library. Library concentrations were determined using the KAPA Library Quantification Kit (Roche) and the average fragment length was determined using a high sensitivity kit with the Bioanalyzer (Agilent). The pooled samples, mock community, and water were sequenced with Illumina MiSeq v2 (500 cycles). Sequences were demultiplexed using oligonucleotide bar code sequences and Illumina software.

Quantitative PCR reactions and raw data processing were conducted using WaferGen SmartChip Real-time PCR system as reported^48^. The qPCR array reactions were performed using 384 previously described and validated primer sets (assays) (Supplementary Table S1)^28^. One sample from M group (2_M) was not included in the qPCR array analysis due to technical error. Samples from pigs that received both mushroom powder and carbadox were also excluded and therefore the results from 16S rRNA amplicon sequencing are not presented in this study.

### 16S sequence analysis and qPCR array data processing

The 16S rRNA amplicon sequences were analyzed using mothur (v 1.39.3)^47^: contigs were made from paired forward and reverse raw reads, aligned to reference sequences (SILVA database release 132)^49^, screened and filtered to remove low quality reads (ambiguous bases allowed = 0, maximum read length = 275, homopolymers allowed = 8), classified with reference to known taxonomic classifications (RDP training set 16)^50^ and clustered into OTUs. The sequences clustered into 137 different OTUs at the 3% dissimilarity level. One NTC group sample (11_NTC) and one AB group sample (16_AB) were discarded due to low number of obtained sequences. 16S sequences were normalized using two different methods: total sum scaling (TSS) using R and rarefying and subsampling using mothur. To produce the rarefied and subsampled OTU table, the data were subsampled to 7,500 reads per sample according to rarefication curves (Supplementary Fig. S4). The rarefied and subsampled OTU table and only quality filtered OTU table (for TSS) were imported into R. After removing the results of samples from animals that received both mushroom powder and carbadox, 132 different OTUs remained in the TSS normalized OTU table and 127 different OTUs in the rarefied and subsampled OTU table. The TSS normalization was carried out in R by dividing each OTU read count by the total number of reads in that sample and all the NA observations (zero sequences) were replaced with 1.490935e-07, which was 100-fold lower than the lowest observed relative abundance.

In the qPCR array, assays “16S old 1_1”,“blaOXY-1_1118”, “cmlV_911”, “czcA_1536”, “fabK_1520”, “intI1F165_clinical_359” and “tetPA_1507” were positive in the negative control, however the Ct-values in the negative control were mostly higher than in experimental samples (Supplementary Table S5). Assay “tetPA_1507” was detected only in the negative control and the Ct-values of assay “czcA_1536” were removed from the results since they were lower in the negative control than in samples. Assay “16S old 1_1” results were removed and not used in normalization since DNA amplification was more efficient in assay “16S new 2_2”. The Ct-values of the remaining four assays that were positive in the negative control were adjusted as follows: The Ct values of each of these assays in each sample were subtracted from Ct value of the assay in the negative control. The resulting numbers were then subtracted from 27, which was the Ct value was used as the cutoff between true positive values and primer-dimer amplification. Next, all the Ct values that were higher than 27 were set to “NA”. After this, all the assays that were undetected in all the samples were removed, resulting in 136 assays out of 382 targeting to AGRs or MGEs being included in the data table. The ΔCt values, ΔΔCt values and relative gene abundances were calculated from these Ct values as previously described^27^. Genes under the detection limit were given a ΔCt value of 20, which was higher than any observed ΔCt (17.4).

### Statistical analyses

R version 3.5.1 (2018-07-02) was used for data exploration, visualization and for all statistical analyses. Analysis of differential abundances of taxa and ARGs and MGEs were carried out using gamma distribution GLMs with TSS normalized OTUs and ARGs and MGEs (relative abundances) and negative binomial GLMs with rarefied and subsampled OTUs (abundance). Gamma distribution model was selected because the relative abundance values (TSS normalized OTUs and ARGs and MGEs) did not fit to normal distribution but followed the gamma distribution. Negative binomial models were selected for rarefied and subsampled OTUs because of the overdispersion in the abundance data. With both model types, p-values were obtained with Tukey’s post-hoc test and adjusted with False discovery rate control^29^ using *glht* function in the multcomp package^51^.

The community and resistome compositions were analyzed using vegan package^52^. PERMANOVA was used to examine the shifts in community composition and resistome between treatment groups with function *adonis* and 9,999 permutations. Nonmetric multidimensional scalings (NMDS) were completed using the Bray-Curtis dissimilarity index with function *metaMDS.* Procrustes analysis with the *protest* function was used to examine the agreement of ordinations of TSS normalized OTUs and rarefied and subsampled OTUs as well as ordination of ARGs and MGEs and both OTU ordinations separately. Mantel’s test and Spearman’s rank correlation was used to analyze the links between microbial community structure and resistome: first, Bray-Curtis dissimilarity indexes were calculated for TSS normalized OTUs, rarefied and subsampled OTUs and for ARGs and MGEs with function *vegdist.* Then the *mantel* function was applied for the dissimilarity indexes of TSS normalized OTUs and ARGs and MGEs as well as rarefied and subsampled OTUs and ARGs and MGEs. Mantel’s tests between both OTU dissimilarity matrices and dissimilarity matrix of ARGs and MGEs were also run for all the treatment groups separately.

To examine if the treatments selected resistance genes into mobile genetic elements, a correlation matrix between ARG and MGE relative abundances was visualized using network analysis with Gephi^53^. Spearman’s rank correlations between ARGs and MGEs within treatment groups and their *p*-values used in network analysis were obtained with package psych^54^ using False discovery rate control^29^. Only ARG-MGE-pairs that were detected at least in three samples with a strong positive correlation (ρ > 0.8, adjusted *p*-value < 0.05) were included.

The factors influencing the shifts in taxonomic structure and resistome were analyzed with machine learning algorithms as previously described^55^. First, dimension reduction was executed on a combined data table of TSS normalized OTUs and ARGs and MGEs using t-SNE algorithm (29) and the R package Rtsne^56^, with 50,000 iterations and “perplexity” set to 5. Then, clusters in the two-dimensional data were identified using HDBSCAN algorithm^31^ in the package dbscan^57^. The minimum number of members in clusters (“minPts”) was set to 3. Classification random forest model^32^ was used with partial dependence plot function in edarf package^58^ for identifying the most important predictors for the clustering pattern. The forests were grown to 10,000 trees using the ranger package^59^ and the best predictors were screened using Gini index by adding predictors one at a time in the order of decreasing importance^60^. The final model was then selected according to the highest Cohen’s Kappa (comparison of observed accuracy and expected accuracy).

## Availability of materials and data

Raw reads from 16S rRNA gene amplicon sequencing are deposited under BioProject accession number PRJNA605462 at NCBI. The R code, mothur commands and all datasets used in statistical analyses are available at https://github.com/sjmuurine/ZnCu.

## Acknowledgements

We thank Ms. Olivia Consoli for excellent technical assistance in laboratory analysis and Purdue University for internal financial support. Information Technology at Purdue is acknowledged for providing computational resources.

## Contributions

J. M. and C. W. analyzed the data (bioinformatics and statistical analyses) and J. M. wrote the manuscript. J. R. was responsible for the practical implementation of the pig feeding trial and fecal sampling. B. R., J. R. and T. A. J. designed the animal experiment and sampling. O.C. performed laboratory work under the supervision of T. A. J. and C. W. All authors contributed to manuscript revisions and have read and approved the final version of the manuscript.

## Corresponding authors

Correspondence to Johanna Muurinen and Timothy A. Johnson, respectively.

## Completing interests statement

The authors declare no competing interests.

## Supplementary information legends

Supplementary Table S1. List of the used primer sets.

Supplementary Fig. S1. Correlation between rarefied and subsampled OTUs and TSS normalized OTUs.

Supplementary Fig. S2 Composition of positive assays grouped by (A) antibiotic group the targeted gene confers resistance, (B) resistance mechanism and (C) mobile genetic element group.

Supplementary Fig. S3. Differentially abundant genera and genes. Samples on x-axis are grouped according to the treatments. Sample names are as follows: NTC= Non-treatment control, AB= Carbadox (antibiotic), M= Mushroom powder and ZnCu= zinc oxide and copper sulfate. The number in front of the group code denotes the number of the pen. Each row represents abundance of each genus or the results of each primer set (assay) (Supplementary Table S1) displayed on the y-axis. Only genera and genes with statistically significant differences between treatment groups are shown (A) Differentially abundant genera with TSS normalization. See Supplementary Table S2 for fold changes. (B) Differentially abundant genera with Rarefying and subsampling. See Supplementary Table S3 for fold changes. (C) Differentially abundant ARGs and MGEs. See Supplementary Table S4 for fold changes.

Supplementary Table S2. Pairwise comparisons of gamma distribution GLMs of relative abundances of each genera between treatment groups. TSS normalized OTU table was used as the input.

Supplementary Table S3. Pairwise comparisons of negative binomial GLMs of abundances of each genera between treatment groups. Rarefied and subsampled OTU table was used as the input.

Supplementary Table S4. Pairwise comparisons of gamma distribution GLMs of relative abundances of each ARG or MGE between treatment groups.

Supplementary Fig. S4. Rarefaction curves. OTU collection curves determined from sequence analysis. Each line represents one sample. Vertical line shows the subsampling cutoff: 7500 sequences

Supplementary Table S5. Assays that had unspecific amplification. Ct values in the negative control and mean Ct-values in samples.

