## Supplemental material for "Swine Growth Promotion with Antibiotics or Alternatives can Increase Antibiotic Resistance Gene Mobility Potential"

Running Title: Growth Promoters Mobilize Antimicrobial Resistance

*Address correspondence to Johanna Muurinen and Timothy Johnson,

, respectively.

Table S1. List of the used primer sets.

| Assay name | Mechanism | Classification | Forward Primer | Reverse Primer |
|---|---|---|---|---|
| 16S old 1_1 | none | housekeeping | GGGTTGCGCTCGTTGC | ATGGYTGTCGTCAGCTCGTG |
| 16S new 2_2 | none | housekeeping | CCTACGGGAGGCAGCAG | ATTACCGCGGCTGCTGGC |
| aacC2_3 | deactivate | Aminoglycoside | ACGGCATTCTCGATTGCTTT | CCGAGCTTCACGTAAGCATTT |
| aacA/aphD_6 | deactivate | Aminoglycoside | AGAGCCTTGGGAAGATGAAGTTT | TTGATCCATACCATAGACTATCTCATCA |
| aac(6)-II_8 | deactivate | Aminoglycoside | CGACCCGACTCCGAACAA | GCACGAATCCTGCCTTCTCA |
| acrB_9 | efflux | MDR | AGTCGGTGTTCGCCGTTAAC | CAAGGAAACGAACGCAATACC |
| acrF_11 | efflux | MDR | GCGGCCAGGCACAAAA | TACGCTCTTCCCACGGTTTC |
| adeA_12 | efflux | MDR | CAGTTCGAGCGCCTATTTCTG | CGCCCTGACCGACCAAT |
| aphA3_14 | deactivate | Aminoglycoside | AAAAGCCCGAAGAGGAACTTG | CATCTTTCACAAAGATGTTGCTGTCT |
| ermK_17 | protection | MLSB | GTTTGATATTGGCATTGTCAGAGAAA | ACCATTGCCGAGTCCACTTT |
| multidrug resistance _20 | efflux | MDR | AATTTTGCCGATTATTGCTGAAA | GATTGTCATCATTCGTTTATCACCAA |
| tet(36)_22 | protection | Tetracycline | AGAATACTCAGCAGAGGTCAGTTCCT | TGGTAGGTCGATAACCCGAAAAT |
| erm(F)_23 | protection | MLSB | CAGCTTTGGTTGAACATTTACGAA | AAATTCCTAAAATCACAACCGACAA |
| cfiA_24 | deactivate | Beta-lactam | GCAGCGTTGCTGGACACA | GTTCGGGATAAACGTGGTGACT |
| Tp614_25 | MGE | Transposase | GGAAATCAACGGCATCCAGTT | CATCCATGCGCTTTTGTCTCT |
| IS613_26 | MGE | Transposase | AGGTTCGGACTCAATGCAACA | TTCAGCACATACCGCCTTGAT |
| blaACC-1_28 | deactivate | Beta-lactam | CACACAGCTGATGGCTTATCTAAAA | AATAAACGCGATGGGTTCCA |
| blaMOX/blaCMY_34 | deactivate | Beta-lactam | CTATGTCAATGTGCCGAAGCA | GGCTTGTCCTCTTTCGAATAGC |
| blaOCH_35 | deactivate | Beta-lactam | GGCGACTTGCGCCGTAT | TTTTCTGCTCGGCCATGAG |
| blaPAO/PDC_36 | deactivate | Beta-lactam | CGCCGTACAACCGGTGAT | GAAGTAATGCGGTTCTCCTTTCA |
| blaVEB_38 | deactivate | Beta-lactam | CCCGATGCAAAGCGTTATG | GAAAGATTCCCTTTATCTATCTCAGACAA |
| bla1_39 | deactivate | Beta-lactam | GCAAGTTGAAGCGAAAGAAAAGA | TACCAGTATCAATCGCATATACACCTAA |
| blaROB_41 | deactivate | Beta-lactam | GCAAAGGCATGACGATTGC | CGCGCTGTTGTCGCTAAA |
| blaOXY-2_42 | deactivate | Beta-lactam | CGTTCAGGCGGCAGGTT | GCCGCGATATAAGATTTGAGAATT |
| blaPSE_43 | deactivate | Beta-lactam | TTGTGACCTATTCCCCTGTAATAGAA | TGCGAAGCACGCATCATC |
| cphA_46 | deactivate | Beta-lactam | GCGAGCTGCACAAGCTGAT | CGGCCCAGTCGCTCTTC |
| bla-L1_48 | deactivate | Beta-lactam | CACCGGGTTACCAGCTGAAG | GCGAAGCTGCGCTTGTAGTC |
| sat4_49 | deactivate | Aminoglycoside | GAATGGGCAAAGCATAAAAACTTG | CCGATTTTGAAACCACAATTATGATA |
| catB3_51 | catB3 | Amphenicol | GCACTCGATGCCTTCCAAAA | AGAGCCGATCCAAACGTCAT |
| catB8_52 | catB8 | Amphenicol | CACTCGACGCCTTCCAAAG | CCGAGCCTATCCAGACATCATT |
| ceoA_53 | efflux | Amphenicol | ATCAACACGGACCAGGACAAG | GGAAAGTCCGCTCACGATGA |
| tet(32)_54 | protection | Tetracycline | CCATTACTTCGGACAACGGTAGA | CAATCTCTGTGAGGGCATTTAACA |
| cmr_57 | efflux | MDR | CGGCATCGTCAGTGGAATT | CGGTTCCGAAAAAGATGGAA |
| dfrA1_58 | protection | Trimethoprim | GGAATGGCCCTGATATTCCA | AGTCTTGCGTCCAACCAACAG |
| dfrA12_59 | protection | Trimethoprim | CCTCTACCGAACCGTCACACA | GCGACAGCGTTGAAACAACTAC |
| acrA_62 | efflux | MDR | GGTCTATCACCCTACGCGCTATC | GCGCGCACGAACATACC |
| emrD_64 | efflux | MDR | CTCAGCAGTATGGTGGTAAGCATT | ACCAGGCGCCGAAGAAC |
| tetU_69 | unknown | Tetracycline | GTGGCAAAGCAACGGATTG | TGCGGGCTTGCAAAACTATC |
| vanC_71 | protection | Vancomycin | CCTGCCACAATCGATCGTT | CGGCTTCATTCGGCTTGATA |
| lmrA_77 | efflux | MLSB | TTCAGATGCAATGGCGTTTG | ATAATCGGGAACATAATGAGCATAACTAC |
| nisB_79 | deactivate | other | GGGAGAGTTGCCGATGTTGTA | AGCCACTCGTTAAAGGGCAAT |
| mdtE/yhiU_81 | efflux | MDR | CGTCGGCGCACTCGTT | TCCAGACGTTGTACGGTAACCA |
| mexA_89 | efflux | MDR | AGGACAACGCTATGCAACGAA | CCGGAAAGGGCCGAAAT |

| | | | | |
|---|---|---|---|---|
| erm(36)_91 | protection | MLSB | GGCGGACCGACTTGCAT | TCTGCGTTGACGACGGTTAC |
| aac(6)-Ib_95 | deactivate | Aminoglycoside | CGTCGCCGAGCAACTTG | CGGTACCTTGCCTCTCAAACC |
| aadA2_97 | deactivate | Aminoglycoside | ACGGCTCCGCAGTGGAT | GGCCACAGTAACCAACAAATCA |
| aadA5_98 | deactivate | Aminoglycoside | ATCACGATCTTGCGATTTTGCT | CTGCGGATGGGCCTAGAAG |
| aadA_99_99 | deactivate | Aminoglycoside | GTTGTGCACGACGACATCATT | GGCTCGAAGATACCTGCAAGAA |
| acrR_102 | regulator | MDR | GCGCTGGAGACACGACAAC | GCCTTGCTGCGAGAACAAA |
| aph(2)-Id_104 | deactivate | Aminoglycoside | TGAGCAGTATCATAAGTTGAGTGAAAAG | GACAGAACAATCAATCTCTATGGAATG |
| cfxA_106 | deactivate | Beta-lactam | TCATTCCTCGTTCAAGTTTTCAGA | TGCAGCACCAAGAGGAGATGT |
| cepA_107 | deactivate | Beta-lactam | AGTTGCGCAGAACAGTCCTCTT | TCGTATCTTGCCCGTCGATAAT |
| blaCMY_108 | deactivate | Beta-lactam | AAAGCCTCATGGGTGCATAAA | ATAGCTTTTGTTTGCCAGCATCA |
| ampC/blaDHA_112 | deactivate | Beta-lactam | TGGCCGCAGCAGAAAGA | CCGTTTTATGCACCCAGGAA |
| blaGES_120 | deactivate | Beta-lactam | GCAATGTGCTCAACGTTCAAG | GTGCCTGAGTCAATTCTTTCAAAG |
| blaSFO_121 | deactivate | Beta-lactam | CCGCCGCCATCCAGTA | GGGCCGCCAAGATGCT |
| blaTLA_122 | deactivate | Beta-lactam | ACACTTTGCCATTGCTGTTTATGT | TGCAAATTTCGGCAATAATCTTT |
| blaZ_123 | deactivate | Beta-lactam | GGAGATAAAGTAACAAATCCAGTTAGATATG | TGCTTAATTTTCCATTTGCGATAAG |
| qacF/H_126 | efflux | MDR | TCGCAACATCCGCATTAAAA | ATGGATTTCAGAACCAGAGAAAGAAA |
| cmlA1_127 | efflux | Amphenicol | TAGGAAGCATCGGAACGTTGAT | CAGACCGAGCACGACTGTTG |
| cmx(A)_129 | efflux | Amphenicol | GCGATCGCCATCCTCTGT | TCGACACGGAGCCTTGGT |
| catA1_130 | deactivate | Amphenicol | GGGTGAGTTTCACCAGTTTTGATT | CACCTTGTCGCCTTGCGTATA |
| sul2_133 | protection | Sulfonamide | TCATCTGCCAAACTCGTCGTTA | GTCAAAGAACGCCGCAATGT |
| ermT_137 | protection | MLSB | GTTCACTAGCACTATTTTTAATGACAGAAGT | GAAGGGTGTCTTTTTAATACAATTAACGA |
| msr(C)_138 | protection | MLSB | TCAGACCGGATCGGTTGTC | CCTATTTTTTGGAGTCTTCTCTCTAATGTT |
| mphB_141 | deactivate | MLSB | CGCAGCGCTTGATCTTGTAG | TTACTGCATCCATACGCTGCTT |
| blaVIM_147 | deactivate | Beta-lactam | GCACTTCTCGCGGAGATTG | CGACGGTGATGCGTACGTT |
| msr(A)_149 | efflux | MLSB | CTGCTAACACAAGTACGATTCCAAAT | TCAAGTAAAGTTGTCTTACCTACACCATT |
| aadD_151 | deactivate | Aminoglycoside | CCGACAACATTTCTACCATCCTT | ACCGAAGCGCTCGTCGTATA |
| nimE_152 | deactivate | MDR | TGCGCCAAGATAGGGCATA | GTCGTGAATTCGGCAGGTTTA |
| Pbp5_153 | protection | Beta-lactam | GGCGAACTTCTAATTAATCCTATCCA | CGCCGATGACATTCTTCTTATCTT |
| pbp_154 | protection | Beta-lactam | CCGGTGCCATTGGTTTAGA | AAAATAGCCGCCCCAAGATT |
| mecA_155 | protection | Beta-lactam | GGTTACGGACAAGGTGAAATACTGAT | TGTCTTTTAATAAGTGAGGTGCGTTAATA |
| emrB/qacA_156 | efflux | MDR | CTTTTCTCTAACCGTACATTATCTACGATAA | AGAACGTAGCGACTGATAAAATGCT |
| blaCTX-M_162 | deactivate | Beta-lactam | GCGATAACGTGGCGATGAAT | GTCGAGACGGAACGTTTCGT |
| aadA2_167 | deactivate | Aminoglycoside | CAATGACATTCTTGCGGGTATC | GACCTACCAAGGCAACGCTATG |
| aadA9_168 | deactivate | Aminoglycoside | CGCGGCAAGCCTATCTTG | CAAATCAGCGACCGCAGACT |
| aphA1_170 | deactivate | Aminoglycoside | TGAACAAGTCTGGAAAGAAATGCA | CCTATTAATTTCCCCTCGTCAAAAA |
| aadE_174 | deactivate | Aminoglycoside | TACCTTATTGCCCTTGGAAGAGTTA | GGAACTATGTCCCTTTTAATTCTACAATCT |
| str_175 | deactivate | Aminoglycoside | AATGAGTTTTGGAGTGTCTCAACGTA | AATCAAAACCCCTATTAAAGCCAAT |
| strA_176 | deactivate | Aminoglycoside | CCGGTGGCATTTGAGAAAAA | GTGGCTCAACCTGCGAAAAG |
| strB_177 | protection | Sulfonamide | GCTCGGTCGTGAGAACAATCT | CAATTTCGGTCGCCTGGTAGT |
| tetA_180 | efflux | Tetracycline | CTCACCAGCCTGACCTCGAT | CACGTTGTTATAGAAGCCGCATAG |
| tetB_181 | efflux | Tetracycline | AGTGCGCTTTGGATGCTGTA | AGCCCCAGTAGCTCCTGTGA |
| tetK_184 | efflux | Tetracycline | CAGCAGTCATTGGAAAATTATCTGATTATA | CCTTGTACTAACCTACCAAAAATCAAAATA |
| tetQ_185 | protection | Tetracycline | CGCCTCAGAAGTAAGTTCATACACTAAG | TCGTTCATGCGGATATTATCAGAAT |
| tetH_187 | efflux | Tetracycline | TTTGGGTCATCTTACCAGCATTAA | TTGCGCATTATCATCGACAGA |
| tetW_191 | protection | Tetracycline | ATGAACATTCCCACCGTTATCTTT | ATATCGGCGGAGAGCTTATCC |

| | | | | |
|---|---|---|---|---|
| tetO_192 | protection | Tetracycline | CAACATTAACGGAAAGTTTATTGTATACCA | TTGACGCTCCAAATTCATTGTATC |
| tetL_195 | efflux | Tetracycline | ATGGTTGTAGTTGCGCGCTATAT | ATCGCTGGACCGACTCCTT |
| tetX_196 | deactivate | Tetracycline | AAATTTGTTACCGACACGGAAGTT | CATAGCTGAAAAAATCCAGGACAGTT |
| tetC_199 | efflux | Tetracycline | ACTGGTAAGGTAAACGCCATTGTC | ATGCATAAACCAGCCATTGAGTAAG |
| tetS_200 | protection | Tetracycline | TTAAGGACAAACTTTCTGACGACATC | TGTCTCCCATTGTTCTGGTTCA |
| tnpA_201 | MGE | Transposase | GCCGCACTGTCGATTTTTATC | GCGGGATCTGCCACTTCTT |
| tnpA_202 | MGE | Transposase | CCGATCACGGAAAGCTCAAG | GGCTCGCATGACTTCGAATC |
| tnpA_203 | MGE | Transposase | GGGCGGGTCGATTGAAA | GTGGGCGGGATCTGCTT |
| tnpA_204 | MGE | Transposase | CATCATCGGACGGACAGAATT | GTCGGAGATGTGGGTGTAGAAAGT |
| tnpA_205 | MGE | Transposase | GAAACCGATGCTACAATATCCAATTT | CAGCACCGTTTGCAGTGTAAG |
| tnpA_206 | MGE | Transposase | TGCAGATGGTTTAACCTTGGATATTT | TCGGTTCATCAAACTGCTTCAC |
| tnpA_207 | MGE | Transposase | AATTGATGCGGACGGCTTAA | TCACCAAACTGTTTATGGAGTCGTT |
| folA_208 | protection | Sulfonamide | CGAGCAGTTCCTGCCAAAG | CCCAGTCATCCGGTTCATAATC |
| ermX_209 | protection | MLSB | GCTCAGTGGTCCCCATGGT | ATCCCCCCGTCAACGTTT |
| VanB_211 | protection | Vancomycin | TTGTCGGCGAAGTGGATCA | AGCCTTTTTCCGGCTCGTT |
| vanD_213 | protection | Vancomycin | CAGAGGAACATAATGTTTCGATAAAATCT | GCCGGATTTTGTGATTCCAA |
| vanHD_214 | protection | Vancomycin | GTGGCCGATTATACCGTCATG | CGCAGGTCATTCAGGCAAT |
| vanHB_215 | protection | Vancomycin | GAGGTTTCCGAGGCGACAA | CTCTCGGCGGCAGTCGTAT |
| vanRA_216 | protection | Vancomycin | CCCTTACTCCCACCGAGTTTT | TTCGTCGCCCCATATCTCAT |
| vanSA_218 | protection | Vancomycin | CGCGTCATGCTTTCAAAATTC | TCCGCAGAAAGCTCAATTTGTT |
| vanWB_220 | protection | Vancomycin | CGGACAAAGATACCCCCTATAAAG | AAATAGTAAATTGCTCATCTGGCACAT |
| vanXB_223 | protection | Vancomycin | AGGCACAAAATCGAAGATGCTT | GGGTATGGCTCATCAATCAACTT |
| vgaB_227 | efflux | MLSB | TAAAAGAGAATAAGGCGCAAGGA | TGTTTAGTAGCATGTTGCATTTTCC |
| pica_229 | protection | MLSB | GCAATCGAGGCGGTGTTC | TTGCCGCAGCCAATTCA |
| mtrE_231 | efflux | MDR | CGATGTGTCGTTTTGGAAGGT | CCTGCACCATGATTCCTCAATA |
| oprD_234 | efflux | MDR | ATGAAGTGGAGCGCCATTG | GGCCACGGCGAACTGA |
| penA_236 | protection | Beta-lactam | AGACGGTAACGTATAACTTTTTGAAAGA | GCGTGTAGCCGGCAATG |
| pmrA_239 | deactivate | Other | TTTGCAGGTTTTGTTCCTAATGC | GCAGAGCCTGATTTCTCCTTTG |
| ttgA_243 | efflux | MDR | ACGCCAATGCCAAACGATT | GTCACGGCGCAGCTTGA |
| ttgB_244 | efflux | MDR | TCGCCCTGGATGTACACCTT | ACCATTGCCGACATCAACAAC |
| mepA_245 | efflux | MDR | ATCGGTCGCTCTTCGTTCAC | ATAAATAGGATCGAGCTGCTGGAT |
| mexE_246 | efflux | MDR | GGTCAGCACCGACAAGGTCTAC | AGCTCGACGTACTTGAGGAACAC |
| qnrA_248 | protection | Fluoroquinolone | AGGATTTCTCACGCCAGGATT | CCGCTTTCAATGAAACTGCAA |
| lnuA_251 | deactivate | MLSB | TGACGCTCAACACACTCAAAAA | TTCATGCTTAAGTTCCATACGTGAA |
| mtrD_253 | efflux | MDR | CGGAGTCCATCGACCATTTG | ATCGTCGGCAAGGAGAATCA |
| vat(E)_263 | deactivate | MLSB | GACCGTCCTACCAGGCGTAA | TTGGATTGCCACCGACAATT |
| ermY_270 | protection | MLSB | TTGTCTTTGAAAGTGAAGCAACAGT | TAACGCTAGAGAACGATTTGTATTGAG |
| cfr_277 | protection | MLSB | GCAAAATTCAGAGCAAGTTACGAA | AAAATGACTCCCAACCTGCTTTAT |
| sulA/folP_280 | protection | Sulfonamide | CAGGCTCGTAAATTGATAGCAGAAG | CTTTCCTTGCGAATCGCTTT |
| ermA/ermTR_283 | protection | MLSB | ACATTTTACCAAGGAACTTGTGGAA | GTGGCATGACATAAACCTTCATCA |
| oleC_285 | efflux | MLSB | CCCGGAGTCGATGTTCGA | GCCGAAGACGTACACGAACAG |
| carB_288 | efflux | MLSB | GGAGTGAGGCTGACCGTAGAAG | ATCGGCGAAACGCACAAA |
| pikR2_290 | protection | MLSB | TCGTGGGCCAGGTGAAGA | TTCCCCTTGCCGGTGAA |
| tetE_291 | efflux | Tetracycline | TTGGCGCTGTATGCAATGAT | CGACGACCTATGCGATCTGA |
| tetbP_294 | efflux | Tetracycline | TGGGCGACAGTAGGCTTAGAA | TGACCCTACTGAAACATTAGAAATATACCT |

| | | | | |
|---|---|---|---|---|
| tetT_297 | protection | Tetracycline | CCATATAGAGGTTCCACCAAATCC | TGACCCTATTGGTAGTGGTTCTATTG |
| tolC_298 | efflux | MDR | GGCCGAGAACCTGATGCA | AGACTTACGCAATTCCGGGTTA |
| vanRB_306 | protection | Vancomycin | GCCCTGTCGGATGACGAA | TTACATAGTCGTCTGCCTCTGCAT |
| vanRC_307 | protection | Vancomycin | TGCGGGAAAAACTGAACGA | CCCCCCATACGGTTTTGATTA |
| vanRC4_308 | protection | Vancomycin | AGTGCTTTGGCTTATCTCGAAAA | TCCGGCAGCATCACATCTAA |
| vanRD_309 | protection | Vancomycin | TTATAATGGCAAGGATGCACTAAAGT | CGTCTACATCCGGAAGCATGA |
| vanSC_311 | protection | Vancomycin | ATCAACTGCGGGAGAAAAGTCT | TCCGCTGTTCCGCTTCTT |
| vanTE_314 | protection | Vancomycin | GTGGTGCCAAGGAAGTTGCT | CGTAGCCACCGCAAAAAAAT |
| vanTC_315 | protection | Vancomycin | ACAGTTGCCGCTGGTGAAG | CGTGGCTGGTCGATCAAAA |
| vanTG_316 | protection | Vancomycin | CGTGTAGCCGTTCCGTTCTT | CGGCATTACAGGTATATCTGGAAA |
| vanYB_317 | protection | Vancomycin | GGCTAAAGCGGAAGCAGAAA | GATATCCACAGCAAGACCAAGCT |
| vanYD_318 | protection | Vancomycin | AAGGCGATACCCTGACTGTCA | ATTGCCGGACGGAAGCA |
| imp-marko_324 | deactivate | Beta-lactam | GGAATAGAGTGGCTTAATTC | GGTTTAACAAAACAACCACC |
| qnrB-bob_resign_328-n | efflux | Amphenicol | GCGACGTTCAGTGGTTCAGA | GCTGCTCGCCAGTCGAA |
| merA-marko_331 | other | MDR | GTGCCGTCCAAGATCATG | GGTGGAAGTCCAGTAGGGTGA |
| int1-a-marko_336 | MGE | Integrase | CGAAGTCGAGGCATTTCTGTC | GCCTTCCAGAAAACCGAGGA |
| intI2_338 | MGE | Integrase | TGCTTTTCCCACCCTTACC | GACGGCTACCCTCTGTTATCTC |
| IncN_rep_340 | MGE | Plasmid-inc | AGTTCACCACCTACTCGCTCCG | CAAGTTCTTCTGTTGGGATTCCG |
| IncN_oriT_341 | MGE | Plasmid-inc | TTGGGCTTCATAGTACCC | GTGTGATAGCGTGATTTATGC |
| IncP_oriT_342 | MGE | Plasmid-inc | CAGCCTCGCAGAGCAGGAT | CAGCCGGGCAGGATAGGTGAAGT |
| IncQ_oriT_343 | MGE | Plasmid-inc | TTCGCGCTCGTTGTTCTTCGAGC | GCCGTTAGGCCAGTTTCTCG |
| IncW_trwAB_344 | MGE | Plasmid-inc | AGCGTATGAAGCCCGTGAAGGG | AAAGATAAGCGGCAGGACAATAACG |
| qacH_351_351 | efflux | MDR | GTCGGTGTTGCTTATGCAGTCT | CAACCAGGCAATGGCTGTAA |
| marR_355 | regulator | MDR | GCTGTTGATGACATTGCTCACA | CGGCGTACTGGTGAAGCTAAC |
| trfa_358 | MGE | Transposase | ACGAAGAAATGGTTGTCCTGTTC | CGTCAGCTTGCGGTACTTCTC |
| intI1F165_clinical_359 | MGE | Integrase | CGAACGAGTGGCGGAGGGTG | TACCCGAGAGCTTGGCACCCA |
| NDM new_362-n2-25-15 | deactivate | Beta-lactam | GGCCACACCAGTGACAATATCA | CAGGCAGCCACCAAAAGC |
| sul1 NEW_363 | protection | Sulfonamide | GCCGATGAGATCAGACGTATTG | CGCATAGCGCTGGGTTTC |
| orf37-IS26_365 | MGE | Insertional | GCCGGGTTGTGCAAATAGAC | TGGCAATCTGTCGCTGCTG |
| orf39-IS26_366 | MGE | Insertional | GCGCGTCGAGCATCAATAG | CAGTTGTGCTGCTGGTGGTC |
| ISPps1-pseud_369 | MGE | Insertional | CACACTGCAAAAACGCATCCT | TGTCTTTGGCGTCACAGTTCTC |
| ISSm2-Xanthob_370 | MGE | Insertional | TGGATCGACCGGTTCCAT | GCTGACCGAGCTGTCCATGT |
| ISAba3-Acineto_371 | MGE | Insertional | TCAGAGGCAGCGGTATACGA | GGTTGATTCAGTTAAAGTACGTAAAACTTT |
| ISEfm1-Entero_372 | MGE | Insertional | AGGTGTCCATGACGTGAAAGTG | TCCTTTGTCCCCTAGGATATTGG |
| mexB_374 | efflux | MDR | CTGGAGATCGACGACGAGAAG | GAAATCGTTGACGTAGCTGGAA |
| cmlA5_375 | efflux | Amphenicol | GCGCTCTTCGAGGATTCG | CCGCCCAAGCAGAAGTAGAC |
| IS1111_376 | MGE | Insertional | GTCTTAAGGTGGGCTGCGTG | CCCCGAATCTCATTGATCAGC |
| PAMBL-1-F_377old_377 | MGE | Plasmid-rep | CAGGCTCTTAATGTGATA | TTATGCTCAATACTCGTG |
| pAKD1-IncP-1__380 | MGE | Plasmid-rep | GGTAAGATTACCGATAAACT | GTTCGTGAAGAAGATGTA |
| pBS228-IncP-1__381 | MGE | Plasmid-rep | CAATCCATCGACAATCAC | GACAATCAGCTACTTCAC |
| IS1133_383 | MGE | Insertional | GCAGCGTCGGGTTGGA | ACGCGTTCGAACAACTGTAATG |
| TN5_384 | MGE | Insertional | CAGCATAAAAAATCCCGACAACA | CCCCGCAACAGACATACGT |
| aac3ia_400 | deactivate | Aminoglycoside | ACGTTCTGCCAAAGTTTGAG | ACTGCCGGATCGTCAC |
| aph4ib_402 | deactivate | Aminoglycoside | GGGAACACCGTGCTCACC | GTTGGTCCCGTGCAGGTC |
| aph3via_403 | deactivate | Aminoglycoside | TCTCATGGCGATATCACGGATAG | TTTCCTCCGATGCATCCTCTC |

| | | | | |
|---|---|---|---|---|
| aph6ic_404 | deactivate | Aminoglycoside | CACGACAACGTGCTCGAC | CCGTCTTCGGCGAACCA |
| ArmA_405 | protection | Aminoglycoside | TCTTCGACGAATGAAAGAGTCG | GCTAATGGATTGAAGCCACAACC |
| spcN_406 | deactivate | Aminoglycoside | GCTATGTGCTGGTGGACTGG | GGAACCACTCGACGAACTCG |
| spec_aph_407 | deactivate | Aminoglycoside | GGTGCTGATATGAATGCCTTTGG | CATTGGGCGCATCAATAAATGG |
| aac(3)-ib_408 | deactivate | Aminoglycoside | CAGCGAGACGTTCATCGC | CACGCTTCAGGTGGCTAATC |
| aac(3)-id_ie_409 | deactivate | Aminoglycoside | AGATAGTTATGCCCGCAACAAG | ACGCGCTGCGCCTATA |
| aac(3)-iid_iii_iif_iia_iie_4 | deactivate | Aminoglycoside | CGATGGTCGCGGTTGGTC | TCGGCGTAGTGCAATGCG |
| aac(3)-xa_411 | deactivate | Aminoglycoside | GCAAGCGGTTCGTGACGTA | TCAGGTGCTCCTCGATCCAG |
| Aac6-Aph2_412 | deactivate | Aminoglycoside | CCAAGAGCAATAAGGGCATACCAA | GCCACACTATCATAACCACTACCG |
| aac(6)-ig_413 | deactivate | Aminoglycoside | GCGATGTTAGAAGCCTCAATTCG | CACACTTCGGCCTGTCGAA |
| aac(6)-iic_414 | deactivate | Aminoglycoside | CAGTCTTTGGCTAATCCATCACAG | AACGAACCCGGCCTTCTC |
| aac(6)-ij_415 | deactivate | Aminoglycoside | ATGCCTGTATCTGAATCCCTGATG | GGCAATCGCTTGTTGAGTATCTG |
| aac(6)-im_417 | deactivate | Aminoglycoside | CGTGAGCATTATACAGAGCAATGG | CCATTTCCGTTCGTAGATATTGGC |
| aac(6)-ir_418 | deactivate | Aminoglycoside | GCTATAACGATCAGCAGCAAGC | CGCGATGCATGGCATGAC |
| aac(6)-is_iu_ix_419 | deactivate | Aminoglycoside | AAGCTTACTCTGGCCTGATCATG | TGCCTGAACGTCGATATTCAGG |
| aac(6)-iv_ih_420 | deactivate | Aminoglycoside | TTGGCTTATACCGACACCCA | CCCGTTGCGATACCTGAAC |
| aac(6)-iw_421 | deactivate | Aminoglycoside | TGCGTCAGTTACTTACACGAAC | CCTGATGCATTGCATGACTGA |
| aac(6)-iz_422 | deactivate | Aminoglycoside | TGCGCCATGACTACGTGAAC | GACTGTCCGAAGCCAGTTCG |
| aacA43_423 | deactivate | Aminoglycoside | CTTGGCCTACATTAGATTCAGCTC | GCTCTCAATCTTTGATAGGAGCAG |
| aadA6_424 | deactivate | Aminoglycoside | CCATCGAGCGTCATCTGGAA | CCCGTCTGGCCGGATAAC |
| aadA7_425 | deactivate | Aminoglycoside | CACTCCGCGCCTTGGA | TGTGGCGGGCTCGAAG |
| aadA10_426 | deactivate | Aminoglycoside | ACAGGCACTCAACGTCATCG | CGCGGAGAACTCTGCTTTGA |
| aadA16_427 | deactivate | Aminoglycoside | ACGGTGGCCTGAAGCC | GAATTGCAGTTCCCGTCTGG |
| aadA17_428 | deactivate | Aminoglycoside | TGTACGGCTCCGCAGTG | CACGGAATGATGTCGTCGTG |
| aadB_429 | deactivate | Aminoglycoside | CCTGCTTGGTGGGCAGAC | CGGCACGCAAGACCTCAA |
| ant4-ib_430 | deactivate | Aminoglycoside | GATGGCCGCTGACACATG | TCAACATTGCGCCATAGTGG |
| ant6-ia_431 | deactivate | Aminoglycoside | TCGCCATGAGCTGCTGA | CCTATCATACTCCGGATAGGCATA |
| aph3-ib_432 | deactivate | Aminoglycoside | AACAGGTTTGGGAGGCGATG | CGCAACAAGCCTCTCCTGAA |
| aph3-viia_433 | deactivate | Aminoglycoside | CTCTCTCATGGAGATATGAGCGCTA | AATCCGGTTCAAGTCCCAACATG |
| aph4-ia_434 | deactivate | Aminoglycoside | CGCTCCCGATTCCGGAA | CACAGTTTGCCAGTGATACACA |
| aph(3)-ia_435 | deactivate | Aminoglycoside | TAACAGCGATCGCGTATTTCG | TCCGACTCGTCCAACATCAATA |
| apmA_436 | deactivate | Aminoglycoside | GGCGCACATGCATTCATCA | CTATACTCCAGTCCCACCATTTGA |
| aph_viii_437 | deactivate | Aminoglycoside | TCGGTATCCCGGTTGTGAG | ACACGAGGTACGGGAATCC |
| acc3-iva_438 | deactivate | Aminoglycoside | CCAACACGACGCTGCATC | GCTGTCGCCACAATGTCG |
| tet40_500 | efflux | Tetracycline | CTGTCCGTGCGCAATATATCC | GGATATATTGCGCACGGACAG |
| tetD_501 | efflux | Tetracycline | AATTGCACTGCCTGCATTGC{EndPos:952} | GACAGATTGCCAGCAGCAGA{EndPos:112 |
| tetPB_502 | efflux | Tetracycline | TGGCAAGACGAGTTTGACTGA | GATCGCTCCACTTCAGCGATAA |
| tet39_505 | efflux | Tetracycline | TATAGCGGGTCCGGTAATAGGTG | CCATAACGATCCTGCCCATAGATAAC |
| tetG_F_507 | efflux | Tetracycline | TCGCGTTCCTGCTTGCC | CCGCGAGCGACAAACCA |
| tetR_506, 508 | regulator | Tetracycline | CCGTCAATGCGCTGATGAC | GCCAATCCATCGACAATCACC |
| dfra14_600 | protection | Trimethoprim | CGGATCATGTCATTGTTTCAGG | ATGTTAGAGGCGAAGTCTTGG |
| dfra17_601 | protection | Trimethoprim | CGGGAACGGCCCTGATATTCC | CGTGTTGCGACCGCATACTTTC |
| dfra7_602 | protection | Trimethoprim | GTAATCGGTAGTGGTCCTGA | ATCAGGACCACTACCGATTAC |
| dfra21_603 | protection | Trimethoprim | TTGTTTCAACGCTGTCGCA | GGTTTCGGTTGAGACAAGCTC |
| dfra5_604 | protection | Trimethoprim | CCATGGAGTGCCAAAGGTG | CACCTTTGGCACTCCATGG |

| | | | | |
|---|---|---|---|---|
| dfrA8_606 | protection | Trimethoprim | GGTCGCACCTGCATCGTTA | AGCGCCACCAATGACGTAG |
| dfrA10_607 | protection | Trimethoprim | CTTCAACTATCACAGAGCACGAAG | TCTACCGGTACATACACATCAGC |
| dfrA15_608 | protection | Trimethoprim | AGGCCGAAAGACTTTCGAGTC | TCACCTTCTGGCTCAATGTCG |
| dfrA18_609 | protection | Trimethoprim | GGAGCGAATCAAGGAGAAAGGAA | GCAATGCGTTGATCGGTATTCTC |
| dfrA22_610 | protection | Trimethoprim | CAGCCGAACACGGCAAAG | CGGAGTGCGTGTACGTGA |
| dfrA25_611 | protection | Trimethoprim | TCAAACTGGACAGCGGCTA | GTCGATTGTCGACACATGCA |
| dfrA27_612 | protection | Trimethoprim | GCCGCTCAGGATCGGTA | GTCGAGATATGTAGCGTGTCG |
| dfrAB4_613 | protection | Trimethoprim | CGGTTCGCATTCCCATCAAA | CGCAGTCATGGATAAATCTGG |
| dfrC_614 | protection | Trimethoprim | GTCGCTCACGATAAACAAAGAGTC | CCCTTCATGGTGAAATGAAGCTTG |
| dfrG_615 | protection | Trimethoprim | TCAATCGGAAGAGCCTTACCTGA | TGGGCAAATACCTCATTCCATTCC |
| dfrK_616 | protection | Trimethoprim | TGCTGCGATGGATAAGAACAG | CTTCCAGGTAATGCTCTTCCG |
| dfrBmulti_617 | protection | Trimethoprim | ACCAAGGCAGAAGTGAAGTCA | GGTGAGCCTCAGACTCGAC |
| fosb_700 | deactivate | Other | CTTGCAGGCCTATGGATTGC | TCTGTTCTCAAGTGTGCCAGTA |
| fosX_702 | deactivate | Other | AGCTGGTTTGTGGATTTGCA | CCACACCGAGAGCTTTAATCCG |
| Arr2_703 | deactivate | Other | TTGGCGATTGGTGACTTGCTAA | ATCGTCTTCGAACGGTCCTG |
| mcr-1_704 | protection | Other | CACATCGACGGCGTATTCTG | CAACGAGCATACCGACATCG |
| sulIII_705 | protection | Sulfonamide | CGCGCTCAAGGCAGATG | GGGAATGCCATCTGCCTTG |
| ere(A)_801 | deactivate | MLSB | GATAATTCTGCTGGCGCACA | GCAGGCGTGGTCACAAC |
| ere(B)_802 | deactivate | MLSB | TCGTATATGGCGGGCGTAGTA | GGTCCAAGATGGGTGAATGCA |
| erm(A)_803 | protection | MLSB | TCGTTGAGAAGGGATTTGCGA | TTGCATGCTTCAAAGCCTGTC |
| erm(B)_804 | protection | MLSB | GAACACTAGGGTTGTTCTTGCA | CTGGAACATCTGTGGTATGGC |
| erm(D)_805 | protection | MLSB | TTTCCGGACAGCATTTGATGC | TCCACTGCCAATACCTTACCG |
| erm(E)_806 | protection | MLSB | GTCACGCAGCTGGAGTTCG | CGGTGAAGCACAGCTCGAC |
| erm(G)_807 | protection | MLSB | CCCTTGAATTAGTACAGAGGTG | GCAAACTCGTATTCCACGA |
| erm(O)_808 | protection | MLSB | TGATGACGGCTCAGTGG | GTGCACCAGCGCCTGA |
| erm(Q)_809 | protection | MLSB | TGAAAGCCATGCGTCTGAC | TTCAGCTGGCAGCTTAAGC |
| erm(S)_810 | protection | MLSB | GAGTACGCCCGCAAACG | GCGTTCGATCCGGAGGA |
| lnuB_811 | deactivate | MLSB | GGATCGTTTACCAAAGGAGAAGG | AGCATAGCCTTCGTATCAGGAA |
| mphA_812 | deactivate | MLSB | TCAGCGGGATGATCGACTG | GAGGGCGTAGAGGGCGTA |
| vat(A)_813 | deactivate | MLSB | ATGAACGGAGCGAATCATCGG | CCATACCGATCCAAACGTCATTTC |
| erm(34)_814 | protection | MLSB | AAAGCGGTTTACAAGCGTTTCG | GGGTGCTCTAGGGTTGTTTAGTG |
| erm(35)_815 | protection | MLSB | CCTTCAGTCAGAACCGGCAA | GCTGATTTGACAGTTGGTGGTG |
| erm(42)_816 | protection | MLSB | TGTTGAGATTGGGCCTGGA | CTAAGGGTGGGTTCTCACTATCTA |
| erm(F)_817 | protection | MLSB | TCTGATGCCCGAAATGTTCAAG | TGAAGGACAATTGAACCTCCCA |
| lnu(F)_818 | deactivate | MLSB | ATACCGGTCATTTCCACTTGGC | GCATCAGGCTGATGAGGTTCAA |
| lsa(C)_819 | protection | MLSB | AAACGGCGTGAAAGTATCAGG | TTGTGGTGATGTAACGGATGC |
| mef(B)_820 | efflux | MLSB | CCGATAGGCTTACTTGTTGCAG | AGTCCACTTGCGGTTTCATTG |
| msr(D)_823 | efflux | MLSB | GGCAAGCTAGGTGTTGAGC | ATTGCTCAACACCTAGCTTGC |
| msr(E)_824 | efflux | MLSB | CGGCAGATGGTCTGAGCTTAAA | CGCACTCTTCCTGCATAAAGGA |
| vgaA_826 | efflux | MLSB | GGAAGCTATAGAGGCGTTTGAATC | CCGAAGGTTCAATACTCAATCGAC |
| vga(A)LC_827 | protection | MLSB | GTGAAGATGTCTCGGGTACAATTG | GAAATACCAGGATTCCCATGCAC |
| vatB_828 | deactivate | MLSB | GCAATTGTTGCTGCGAATTCAG | GTGCTGACCAATCCCACCA |
| cat_900 | deactivate | Phenicol | ATCGGCCAGACTGGATATCGA | CACAGCTCCAGTTGCAACAAC |
| catA2_902 | deactivate | Phenicol | CCTGGAACCGCAGAGAACA | CGGAACTCCGGAAACTGATTAAC |
| catA3_903 | deactivate | Phenicol | CTGATTGCTCAGGCCGTGAA | ATGAGTATGGGCAACTCAGTGC |

| | | | | |
|---|---|---|---|---|
| catB2_904 | deactivate | Phenicol | GCTACTATTCCGGCTATTACCATG | GGGCTCCTCGTTCATGTAGA |
| catB9_906 | deactivate | Phenicol | CACCTTATGAAGTGGTCGGTTCA | GTCTGATGAACACAGAGACTGCA |
| cat(pC221)_908 | deactivate | Phenicol | AATGACCGTATGCTGCAAGAAG | TTTGCCTGCTATGGCATTCTG |
| catP_909 | deactivate | Phenicol | CCTTTGGACTGAGTGTAAGTCTGA | TAAAGCCATCGAAGGTTGACCA |
| catQ_910 | deactivate | Phenicol | AGGTGCACTTACAGTATGACTGC | AACGTGGGAAGTTCTCGTCATAC |
| cmlV_911 | deactivate | Phenicol | GCCCTCATCACCGTCTTCG | GGACGTTGGCGATGGAGAG |
| fexA_912 | efflux | Phenicol | TGGTGTGGCTGTTGCAATCTTA | CCAAGGTACAAAGCACCTTGGA |
| floR_913 | efflux | Amphenicol | AACCCGCCCTCTGGATCA | GCCGTCGAGAAGAAGACGAA |
| vanG_1002 | protection | Vancomycin | TGTTTCGCAGAACCGTGTCAA | CCCTGCACTGTTCCATCTTCTC |
| vanC2/vanC3_1003 | protection | Vancomycin | TGACTGTCGGTGCTTGTGA | GATAGAGCAGCTGAGCTTGTTC |
| beta_ccra_1104 | deactivate | Beta-lactam | CACTGGCACGGCGATTGTA | CGGCAGCCAAACCACGATA |
| cefa_ampc_1105 | deactivate | Beta-lactam | CAGGATCTGATGTGGGAGAACTA | TCGGGAACCATTTGTTGGC |
| bl1acc_1107 | deactivate | Beta-lactam | TGTTATCCGTGATTACCTGTCTGG | CTCAGCGAGCCAACTTCAAATA |
| blaCTX-M-1,3,15_1108 | deactivate | Beta-lactam | CGTACCGAGCCGACGTTAA | CAACCCAGGAAGCAGGCA |
| blaSHV-11_1110 | deactivate | Beta-lactam | TTGACCGCTGGGAAACGG | TCCGGTCTTATCGGCGATAAAC |
| bl3_cpha_1113 | deactivate | Beta-lactam | GTAACGCCTACTGGAAGTCCA | CAGCTTCTCCTTGAGAATGCAG |
| blaB-11,13,14_1114 | deactivate | Beta-lactam | CGTGCCGGAGGTCTTGAATA | GGGATAGTAAACCTGAAACTCGGA |
| blaIND_1115 | deactivate | Beta-lactam | CGCCTGTTAAACCCAACCTGTA | CGCTCTGTCATCATGAGAGTGG |
| blaLEN_1116 | deactivate | Beta-lactam | TGTTCGCCTGTGTGTTATCTCC | GCAGCACTTTAAAGGTGCTCAC |
| blaOXY-1_1118 | deactivate | Beta-lactam | AAAGGTGACCGCATTCGC | CCAGCGTCAGCTTGCG |
| bla-SME_1119 | deactivate | Beta-lactam | GAGGAAGACTTTGATGGGAGGATTG | CGCTATATTGCAATGCAGCAGAAG |
| blaCARB_1120 | deactivate | Beta-lactam | TGATTTGAGGGATACGACAACTCC | CTGTAATACTCCGAGCACCAA |
| blaGOB_1121 | deactivate | Beta-lactam | CTTGGGCTTGAATGCTCAGGTA | TGTATGGTCGTAGTGAGCCTGA |
| blaHERA_1122 | deactivate | Beta-lactam | GGGCAACCGCATTCTGAC | GCATCTCCCACTTTATCGTCAC |
| blaMIR_1123 | deactivate | Beta-lactam | CGGTCTGCCGTTACAGGTG | AAAGACCCGCGTCGTCATG |
| blaFOXnew_1125 | deactivate | Beta-lactam | CCTACGGCTATTCGAAGGAAGATAAG | CCGGATTGGCCTGGAAGC |
| nonmobile_blaADC_1127 | deactivate | Beta-lactam | GGTATGGCTGTGGGTGTTATTCA | AGGCAAGGTTACCACTTGTATACG |
| nonmobile blaBEL_1128 | deactivate | Beta-lactam | ATGTCCATGGCACAGACTGTG | CCTGTCTTGTCACCCGTTACC |
| blaIMI_1129 | deactivate | Beta-lactam | ACATCTACACCTGCAGCAGTAG | AATCGCTTGGTACGCTAGCA |
| norA_1200 | efflux | Fluoroquinolone | ATCGCCGTTTGGTGGTACG | TCCACCAATCCCTGGTCCTAAA |
| qepA_1_2_1201 | efflux | Fluoroquinolone | GGGCATCGCGCTGTTC | GCGCATCGGTGAAGCC |
| qnrB4_1202 | protection | Fluoroquinolone | TCACCACCCGCACCTG | GGATATCTAAATCGCCCAGTTCC |
| qnrS1_S3_S5_1203 | protection | Fluoroquinolone | CCACTTTGATGTCGCAGATCTTC | CCCTCTCCATATTGGCATAGGAAA |
| qnrVC1_VC3_VC6_1204 | protection | Fluoroquinolone | CTCACATCAGGACTTGCAAGAA | ATGAAGCATCTCGAAGATCAGC |
| qnrVC4_VC5_VC7_1205 | protection | Fluoroquinolone | TTCCTTTAAACGGGCAAACCTC | CGATACCTGATTCATGAAGCTAGC |
| mdth_1300 | efflux | MDR-chromo | ATGCTGGCTGTACAAGTGATG | CACTCCAGCGGGCGATA |
| cefa_qacelta_1301 | efflux | MDR-mobile | TAGTTGGCGAAGTAATCGCAAC | TGCGATGCCATAACCGATTATG |
| mdtg_1302 | efflux | MDR | TTCCAGCCGGTCAGCAA | GACATCTCCCGCGAGTTCG |
| pcoA_1303 | deactivate | MDR-mobile | TGGCGTATGGAGTTTCAATGC | GAATAATGCCGTGCCAGTGAA |
| silE_1304 | deactivate | MDR-mobile | GGTGGAAAGTCATCAGAGGATGA | CAAAGCCCAGCAAGGATGC |
| arsA_1305 | efflux | MDR-mobile | CAGGTCAGCCGCATCAACC | GCCTGAAACACGGCAATTTCTTC |
| qacA/B_1306 | efflux | MDR-mobile | AAGGGCCACTGCATTAGCTG | CCAGTCCAATCATGCCTGCA |
| qacF/H_1308 | efflux | MDR-mobile | CTGAAGTCTAGCCATGGATTCACTAG | CAAGCAATAGCTGCCACAAGC |
| bacA_1500 | deactivate | Other | ATCCGCGGCACCCTGA | CCTGCTTGATGGACTTGATGAAGA |
| aac(6)l1_1501 | deactivate | Aminoglycoside | GGGAATTATCGGAATAGCTCTTGG | TTGGGCTGTTCTTCCTAGCTAA |

| | | | | |
|---|---|---|---|---|
| aac(6)-ly_1502 | deactivate | Aminoglycoside | GCCTCAATCCGCCACGATTA | ACGCGCTCTGTTTCCTCAAA |
| aph6ia_1503 | deactivate | Aminoglycoside | CGCTGGGAGCTGAAGAGG | AGCATCGTGCTGCTCTCC |
| bexA/norM_1504 | efflux | MDR | TCGGGCATCCCGTTTATGATC | GTAGGCTGCGCATAATACCCA |
| ampC_1505 | deactivate | Beta-lactam | CTGGCGCATACCTGGATTAC | GCCAGTTCAGCATCTCCCA |
| blaOXA10_1506 | deactivate | Beta-lactam | CGACCGAGTATGTACCTGCTTC | TCAAGTCCAATACGACGAGCTA |
| tetPA_1507 | efflux | Tetracycline | GGAAACCTTAGTTCAGTGACTTGG | CCCATTTAACCACGCACTGAA |
| blaIMIR_1508 | regulator | Beta-lactam | AGCCGGACTAGAGCTTCATG | GGCAGAACTCATCATCTGCAAA |
| mdtA_1509 | efflux | MDR | ACAAGCCCAGGGCCAAC | CCTTAATGGTGCCTTCGGTTTC |
| aac3-Via_1510 | deactivate | Aminoglycoside | GTGTCCGTCGCCAAGGA | GGTGACGGCCTTGTCGA |
| mefA_1511 | efflux | MLSB | TAATTATCGCAGCAGCTGGTTC | GTTCCCAAACGGAGTATAAGAGTG |
| blaTEM_1512 | deactivate | Beta-lactam | CGCCGCATACACTATTCTCAG | GCTTCATTCAGCTCCGGTTC |
| tetM_1513 | protection | Tetracycline | GGAGCGATTACAGAATTAGGAAGC | TCCATATGTCCTGGCGTGTC |
| vanA_1514 | protection | Vancomycin | GGGCTGTGAGGTCGGTTG | TTCAGTACAATGCGGCCGTTA |
| vanXA_1515 | protection | Vancomycin | TCGTTGGGACGCTAAATATGC | GGACGGTAACCGTCCCATA |
| tetJ_1516 | efflux | Tetracycline | CAGCGCCCATACGCCATTTA | CCTACTTCAGTAGTGTGCCAAGC |
| blaPER_1517 | deactivate | Beta-lactam | GCAAATGAAGCGCAGATGC | GACCACAGTACCAGCTGGTA |
| tet(38)_1518 | efflux | Tetracycline | AAGCGACATTAGCCGGTTTAG | CTGCTCGTACTTAAGCCAAGG |
| lnuC_1519 | deactivate | MLSB | GGGTGTAGATGCTCTTCTTGGA | CTTTACCCGAAAGAGTTTCTACCG |
| fabK_1520 | protection | Other | CAGGAGCAGGAAATCCAAGC | CCAGCTTCCATTCCTTCTGC |
| vanSB_1521 | protection | Vancomycin | GAAGATAAAGAGGGAAGCGTACTC | CCGAATTGTCAGCCCTTGATAA |
| intI3_1522 | MGE | Integrase | CAGGTGCTGGGCATGGA | CCTGGGCAGCATCACCA |
| KPC_1523 | deactivate | Beta-lactam | GCCGCCAATTTGTTGCTGAA | GCCGGTCGTGTTTCCCTTT |
| mobA_1524 | mobA | Plasmid | GCTTCCCGTAACGAGGTAGT | CCTTGAACGGTATCAGCACG |
| traN_1525 | MGE | Plasmid | GCTTGGCGGTCAGCAATT | TTAGGAATAACAATCGCTACACCTTTA |
| tra-A_1526 | MGE | Plasmid | AAGTGTTCAGGGTGCTTCTGCGC | GTCATGTACATGATGACCAAAA |
| trb-C_1527 | MGE | Plasmid | CGGYATWCCGSCSACRCTGCG | GCCACCTGYSBGCAGTCMCC |
| ISCR1_1528 | MGE | Insertional | ATGGTTTCATGCGGGTT | CTGAGGGTGTGAGCGAG |
| copA_1529 | other | MDR-mobile | TGCACCTGACVGGSCAYAT | GVACTTCRCGGAACATRCC |
| Staphylococci_1532 (mec | other | Taxonomic | CGCAACGTTCAATTTAATTTTGTTAA | TGGTCTTTCTGCATTCCTGGA |
| A.baumannii_1535 (ompA | other | Taxonomic | TCTTGGTGGTCACTTGAAGC | ACTCTTGTGGTTGTGGAGCA |
| czcA_1536 | efflux | MDR-mobile | GCCTTGTTCATCGGCGAAC | GGCAATGTCGCCTTCGTTC |
| optrA_1538 | protection | Phenicol | GGTGGATGAAGTCCGTACGG | AGGTTAGACCTCCAAGAGCCA |
| tet44_1539 | protection | Tetracycline | CTCATGTAGATGCAGGAAAGACG | GTAACTGCTGCCTGAATTGTGA |
| aph3-III_1540 | deactivate | Aminoglycoside | CAGAAGGCAATGTCATACCACTTG | GACAGCCGCTTAGCCGAA |
| ant6-ib_1541 | deactivate | Aminoglycoside | AGAACATCCGACAGCACGTTC | CCAACCTTCCATGAAATCATTCGC |
| ARR-3_1542 | deactivate | Other | GATCGTCTTCGAACGGTCCTG | TTTGGCGATTGGTGACTTGCT |
| mcr-2_1543 | protection | Other | CGGCGTACTTTAAGCGTTATGATG | GCATTTGGCATACCATGCAGATAG |
| bla-ACT_1544 | deactivate | Beta-lactam | AAGCCGCTCAAGCTGGA | GCCATATCCTGCACGTTGG |
| aac(3)-Xa_1545 | deactivate | Aminoglycoside | TGTACGGCTCCGCAGTG | CACGGAATGATGTCGTCGTG |
| IS26_1546 | MGE | Insertional | ATGGATGAAACCTACGTGAAGGTC | CGGTACTTAATCTGTCGGTGTTCA |
| IS3_1547 | MGE | Insertional | CGGTCTGAGCTTCGGGAA | AGAACTGTCACTCCGGTCTG |
| IS256_1548 | MGE | Insertional | CTTGCGCATCATTGGATGATGG | AAGAACGGCTCCAATTAAGCGA |
| sugE_1549 | efflux | MDR-mobile | CTTAGTTATTGCTGGTCTGCTGGA | GCATCGGGTTAGCGGACTC |
| ISEcp1_1550 | MGE | Insertional | CATGCTCTGCGGTCACTTC | GACGCACCTTCTTGATGACC |
| IS200_1551 | MGE | Insertional | CCAAATACCGAAGACAAGCGTTC | CCAAACTGCTCGTAAAGCATCAG |

| | | | | |
|---|---|---|---|---|
| IS1247_1552 | MGE | Insertional | CGGCCGTCACTGACCAA | TCGGCAGGTTGGTGACG |
| IS630_1553 | MGE | Insertional | CCGCCACCAGTGTGATGG | TTGGCGCTGACTGGATGC |
| TN5403_1556 | MGE | Transposase | AAGCGAATGGCGCGAAC | CGCGCAGGGTAAACTGC |
| IS200_1557 | MGE | Insertional | GCACACCCGATGGAACTGTAAA | TCGGCGGGATCTCCAGAAG |
| IS21-ISAs29_1558 | MGE | Insertional | GGTCCGTCAGGCACAAGTC | GGGATCGTATCGGCAAGCC |
| Tn3_1559 | MGE | Transposase | GCTGAGGTGTTCAGCTACATCC | GCTGAGGTAGTCACAGGCATTC |
| IS6/257_1560 | MGE | Insertional | ATATCGTGCCATTGATGCAGAG | ACCATTGCTACCTTCGTTGAAG |
| IS6100_1561 | MGE | Insertional | CGCACCGGCTTGATCAGTA | CTGCCACGCTCAATACCGA |
| IS15DI_1562 | MGE | Insertional | CAATACCTTTGATGGTGGCGTAAG | CTTACGCCACCATCAAAGGTATTG |
| IncN_korA_1563 | MGE | Plasmid-inc | GGAACGTTTGTAYCTTGTATTG | ACTCACTATCTTCTGTTGATTG |
| IncF_FIC_1564 | MGE | Plasmid-inc | GTGAACTGGCAGATGAGGAAGG | TTCTCCTCGTCGCCAAACTAGAT |
| IncI1_repI1_1565 | MGE | Plasmid-inc | CGAAAGCCGGACGGCAGAA | TCGTCGTTCCGCCAAGTTCGT |
| IncHI2-smr0018_1566 | MGE | Plasmid-inc | ATAATGATTCACCGGGGTAG | CTTCAGGCTATCGTTTCG |
| IS91_1567 | MGE | Insertional | GGATGCCACTGCTGGTCA | ACAGTGGATACAGTATCTGCTGAG |
| IS5/IS1182_1568 | MGE | Insertional | TTCTCGAAGAATCGCCATGGC | GCTTTGGATCGCTCCAATCGA |
| cro_1569 | MGE | Other | AGATGTTATCGACCACTTCGGA | CCGCTTGGCGATAAGCG |
| EAE_05855_1570 | MGE | Other | CCCATCACCGCTGAACTGG | TGGGCGCTGCCATCTAAAC |
| tcrB_1571 | efflux | MDR-mobile | GTGCCGGAACTCAAGTAGCA | GCACCGACTGCTGGACTTAA |
| terW_1572 | other | MDR-mobile | TCAAAGAGCTACGCGAGTCATA | CCTTCCCTGTGGACTCACC |
| pbrT_1573 | efflux | MDR-mobile | GATGCGCACTGGGCTTG | TCGGAATATGCGGAAATGCG |
| qnrD_1574 | protection | Fluoroquinolone | CGCTGGAATGGCACTGTGA | GCTCTCCATCCAACTTCACTCC |
| cadC_1575 | other | MDR-mobile | CGCTCTGTGTCAGGATGAAGAG | CTTTCTTATGTGCTAGGGCGATCA |
| qnrS2_1576 | protection | Fluoroquinolone | TCCCGAGCAAACTTTGCCAA | GGTGAGTCCCTATCCAGCGA |
| oqxA_1577 | other | Fluoroquinolone | GAGTCAACCTACCTCCACTATCA | GCTGCGAGTTATCCAGCAG |
| adeI_1578 | efflux | MDR | CAGTCTGGTTTGCAGTAACCA | CACTCCTACAACAACAGGCAA |
| qnrB46,47,48_1579 | protection | Fluoroquinolone | CGACGTTCAGTGGTTCAGATCTC | GCCAAGCCGCTCCATGAG |

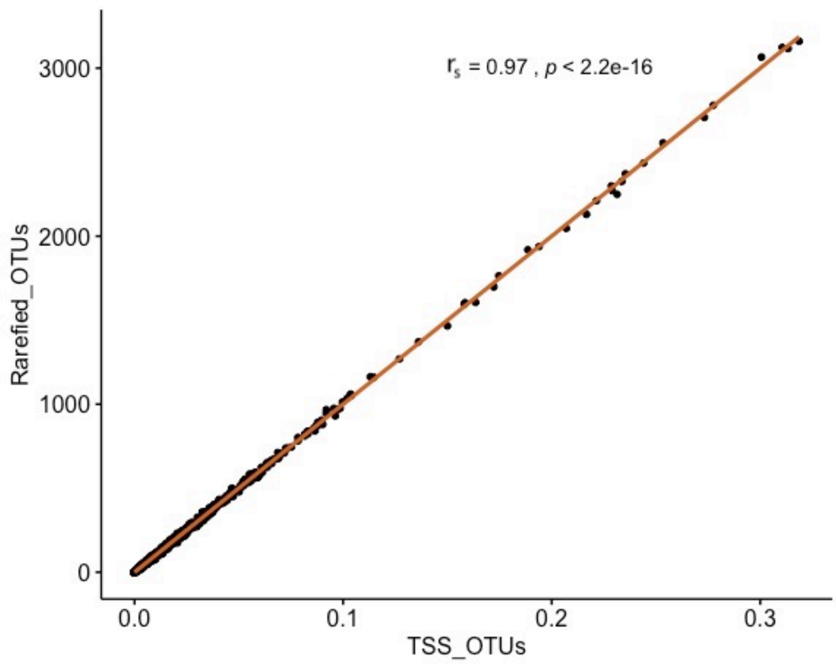


Fig. S1 Correlation between rarefied and subsampled OTUs and TSS normalized OTUs.

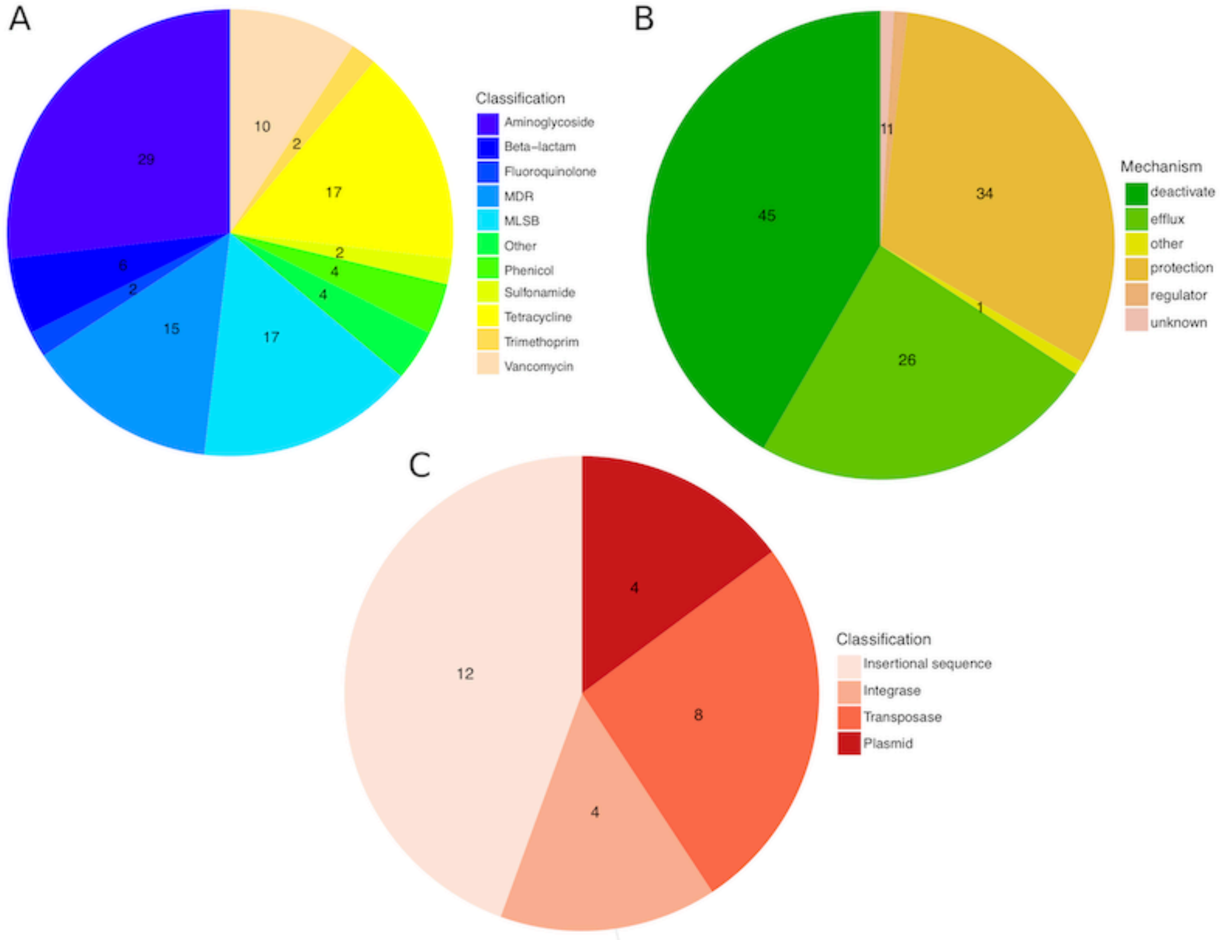


Fig. S2 Composition of positive assays grouped by (A) antibiotic group the targeted gene confers resistance, (B) resistance mechanism and (C) mobile genetic element group.

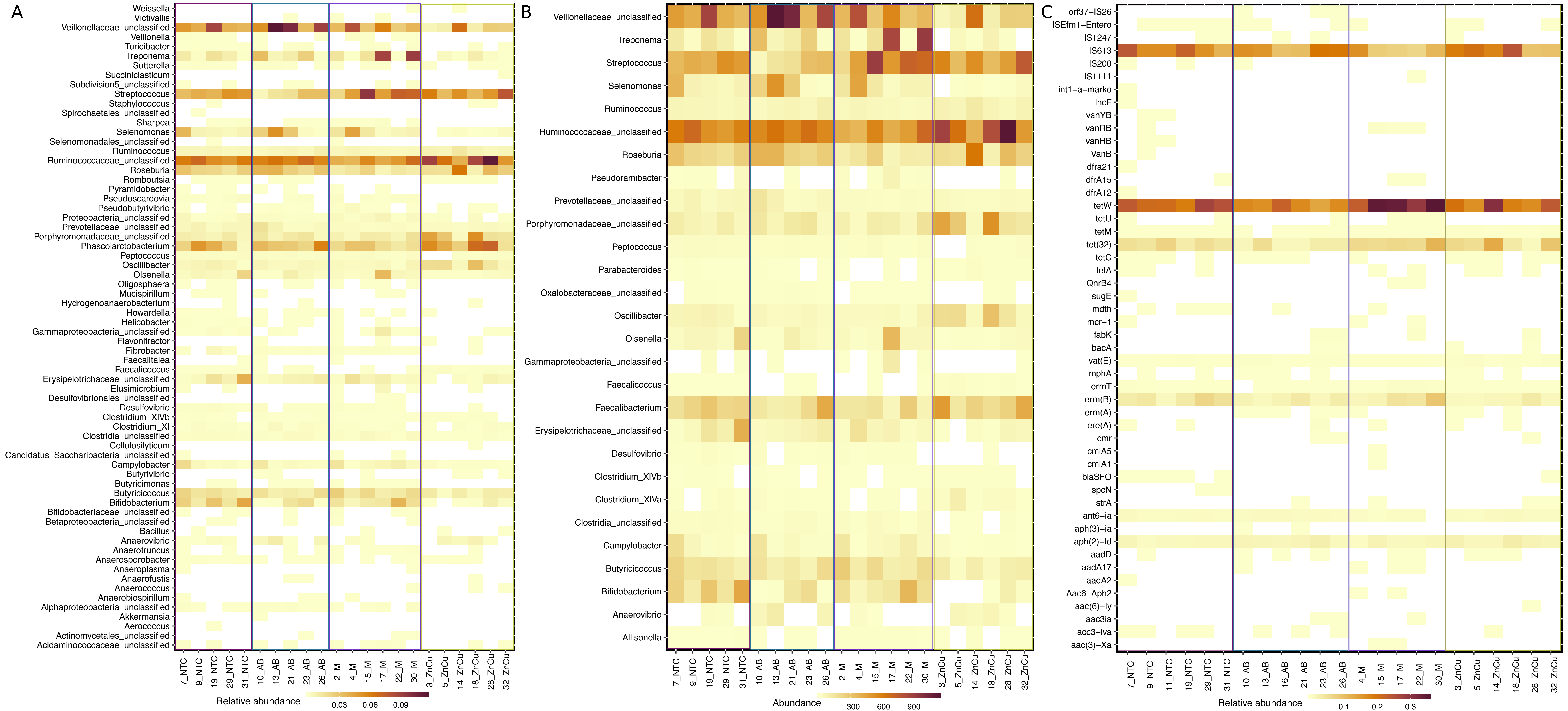

Fig. S3. Differentially abundant genera and genes. Samples on x-axis are grouped according to the treatments. Sample names are as follows: NTC= Non-treatment control, AB= Carbadox (antibiotic), M= Mushroom powder and ZnCu= zinc oxide and copper sulfate. The number in front of the group code denotes the number of the pen. Each row represents abundance of each genus or the results of each primer set (assay) (Supplementary Table S1) displayed on the y-axis. Only genera and genes with statistically significant differences between treatment groups are shown (A) Differentially abundant genera with TSS normalization. See Supplementary Table S2 for fold changes. (B) Differentially abundant genera with Rarefying and subsampling. See Supplementary Table S3 for fold changes. (C) Differentially abundant ARGs and MGEs. See Supplementary Table S4 for fold changes

Table S2. Pairwise comparisons of gamma distribution GLMs of relative abundances of each genera between treatment groups. TSS normalized OTU table was used as the input.

| Comparison ($X_1$ - $X_2$) | Genus | Delta Estimate (X1 - X2) [a] | Std. Error | z-value | p.adjusted | Fold Change (X1 / X2) [b] |
|---|---|---|---|---|---|---|
| AB - ZnCu | Veillonellaceae_unclassified | 1.21 | 0.39 | 3.14 | 9.43E-03 | 3.352 |
| AB - ZnCu | Oscillibacter | -1.32 | 0.33 | -3.99 | 3.75E-04 | 0.267 |
| AB - ZnCu | Porphyromonadaceae_unclassified | -0.98 | 0.34 | -2.93 | 1.79E-02 | 0.374 |
| AB - ZnCu | Streptococcus | -0.93 | 0.25 | -3.69 | 1.28E-03 | 0.396 |
| AB - ZnCu | Selenomonas | 3.04 | 0.68 | 4.46 | 3.67E-05 | 21.000 |
| AB - ZnCu | Clostridia_unclassified | 1.70 | 0.47 | 3.62 | 1.59E-03 | 5.458 |
| AB - ZnCu | Campylobacter | 1.97 | 0.75 | 2.63 | 4.29E-02 | 7.188 |
| AB - ZnCu | Desulfovibrio | 4.34 | 0.90 | 4.84 | 5.51E-06 | 76.588 |
| AB - ZnCu | Subdivision5_unclassified | 5.75 | 0.84 | 6.88 | 1.31E-11 | 313.962 |
| AB - ZnCu | Elusimicrobium | -5.86 | 1.15 | -5.09 | 1.26E-06 | 0.003 |
| AB - ZnCu | Fibrobacter | 4.19 | 0.96 | 4.38 | 8.57E-05 | 66.003 |
| AB - ZnCu | Pseudoscardovia | 7.72 | 0.55 | 13.91 | 0.00E+00 | 2252.752 |
| AB - ZnCu | Alphaproteobacteria_unclassified | 8.00 | 0.70 | 11.46 | 0.00E+00 | 2966.181 |
| AB - ZnCu | Butyricimonas | 5.48 | 0.63 | 8.76 | 0.00E+00 | 240.090 |
| AB - ZnCu | Helicobacter | 6.18 | 0.73 | 8.50 | 0.00E+00 | 485.265 |
| AB - ZnCu | Sharpea | 5.91 | 0.77 | 7.68 | 3.90E-14 | 368.828 |
| AB - ZnCu | Bifidobacteriaceae_unclassified | 3.67 | 0.90 | 4.09 | 2.24E-04 | 39.081 |
| AB - ZnCu | Oligosphaera | 4.87 | 0.64 | 7.58 | 7.14E-14 | 129.672 |
| AB - ZnCu | Veillonella | 5.69 | 0.75 | 7.55 | 9.17E-14 | 295.335 |
| AB - ZnCu | Anaerotruncus | 3.06 | 1.09 | 2.80 | 2.60E-02 | 21.242 |
| AB - ZnCu | Mucispirillum | 5.58 | 0.96 | 5.80 | 6.70E-08 | 264.733 |
| AB - ZnCu | Betaproteobacteria_unclassified | 3.67 | 0.90 | 4.08 | 2.49E-04 | 39.081 |
| AB - ZnCu | Staphylococcus | -4.44 | 0.81 | -5.48 | 1.53E-07 | 0.012 |
| AB - ZnCu | Anaerobiospirillum | 5.40 | 1.08 | 5.01 | 2.36E-06 | 221.168 |
| AB - ZnCu | Anaerococcus | -4.10 | 0.94 | -4.38 | 6.75E-05 | 0.017 |
| AB - ZnCu | Faecalitalea | 3.52 | 1.15 | 3.05 | 1.23E-02 | 33.792 |
| AB - ZnCu | Turicibacter | 2.41 | 0.87 | 2.76 | 2.92E-02 | 11.122 |
| AB - ZnCu | Cellulosilyticum | -4.90 | 0.78 | -6.31 | 1.31E-09 | 0.007 |
| AB - ZnCu | Weissella | -4.72 | 1.08 | -4.36 | 8.08E-05 | 0.009 |
| AB - ZnCu | Actinomycetales_unclassified | 3.83 | 0.80 | 4.81 | 8.08E-06 | 46.196 |
| AB - ZnCu | Victivallis | 3.67 | 0.83 | 4.43 | 6.70E-05 | 39.081 |
| AB - ZnCu | Bacillus | -4.96 | 1.12 | -4.41 | 6.00E-05 | 0.007 |
| AB - ZnCu | Succiniclasticum | -6.25 | 0.78 | -8.01 | 2.22E-15 | 0.002 |
| AB - ZnCu | Pyramidobacter | 5.42 | 0.79 | 6.85 | 8.72E-11 | 224.996 |
| AB - ZnCu | Akkermansia | 5.70 | 0.64 | 8.95 | 0.00E+00 | 297.725 |
| M - AB | Roseburia | -0.86 | 0.29 | -2.93 | 1.74E-02 | 0.425 |
| M - AB | Streptococcus | 1.06 | 0.25 | 4.21 | 1.86E-04 | 2.878 |

| | | | | | | |
|---|---|---|---|---|---|---|
| M - AB | Prevotellaceae_unclassified | -1.08 | 0.34 | -3.19 | 7.44E-03 | 0.340 |
| M - AB | Elusimicrobium | 8.62 | 1.15 | 7.49 | 1.45E-13 | 5531.182 |
| M - AB | Anaerovibrio | -2.73 | 0.90 | -3.02 | 1.32E-02 | 0.065 |
| M - AB | Clostridium_XI | -1.63 | 0.62 | -2.62 | 4.38E-02 | 0.195 |
| M - AB | Helicobacter | -2.15 | 0.73 | -2.95 | 1.69E-02 | 0.117 |
| M - AB | Gammaproteobacteria_unclassified | 3.23 | 1.13 | 2.85 | 2.26E-02 | 25.292 |
| M - AB | Bifidobacteriaceae_unclassified | 2.47 | 0.90 | 2.76 | 2.92E-02 | 11.871 |
| M - AB | Veillonella | 1.94 | 0.75 | 2.58 | 4.85E-02 | 6.982 |
| M - AB | Sutterella | -2.04 | 0.61 | -3.33 | 4.81E-03 | 0.130 |
| M - AB | Selenomonadales_unclassified | 2.91 | 0.97 | 2.99 | 1.48E-02 | 18.286 |
| M - AB | Anaerococcus | 3.43 | 0.94 | 3.66 | 1.39E-03 | 30.910 |
| M - AB | Anaerofustis | -4.43 | 0.87 | -5.11 | 2.40E-06 | 0.012 |
| M - AB | Hydrogenoanaerobacterium | -3.83 | 1.17 | -3.27 | 5.95E-03 | 0.022 |
| M - AB | Desulfovibrionales_unclassified | 4.35 | 0.37 | 11.86 | 0.00E+00 | 77.587 |
| M - AB | Candidatus_Saccharibacteria_unclassified | 4.10 | 0.84 | 4.90 | 3.95E-06 | 60.265 |
| M - AB | Weissella | 3.43 | 1.08 | 3.17 | 8.20E-03 | 30.910 |
| M - AB | Romboutsia | -5.31 | 0.50 | -10.59 | 0.00E+00 | 0.005 |
| M - AB | Bacillus | 3.44 | 1.12 | 3.05 | 1.20E-02 | 31.039 |
| M - AB | Anaeroplasma | 4.51 | 1.00 | 4.52 | 4.88E-05 | 91.064 |
| M - AB | Akkermansia | -5.70 | 0.64 | -8.95 | 0.00E+00 | 0.003 |
| M - NTC | Butyricimonas | 1.65 | 0.63 | 2.64 | 4.05E-02 | 5.224 |
| M - NTC | Helicobacter | -3.20 | 0.73 | -4.39 | 9.84E-05 | 0.041 |
| M - NTC | Sutterella | -1.64 | 0.61 | -2.67 | 3.78E-02 | 0.194 |
| M - NTC | Staphylococcus | -4.74 | 0.81 | -5.86 | 1.45E-08 | 0.009 |
| M - NTC | Anaerococcus | 3.43 | 0.94 | 3.66 | 1.53E-03 | 30.910 |
| M - NTC | Flavonifractor | 4.40 | 1.01 | 4.35 | 7.67E-05 | 81.446 |
| M - NTC | Hydrogenoanaerobacterium | -3.67 | 1.17 | -3.13 | 9.69E-03 | 0.026 |
| M - NTC | Desulfovibrionales_unclassified | 4.35 | 0.37 | 11.86 | 0.00E+00 | 77.587 |
| M - NTC | Butyrivibrio | 3.44 | 1.09 | 3.14 | 8.73E-03 | 31.039 |
| M - NTC | Weissella | 3.43 | 1.08 | 3.17 | 8.66E-03 | 30.910 |
| M - NTC | Actinomycetales_unclassified | 3.95 | 0.80 | 4.95 | 3.51E-06 | 51.817 |
| M - NTC | Romboutsia | -5.58 | 0.50 | -11.13 | 0.00E+00 | 0.004 |
| M - NTC | Spirochaetales_unclassified | -4.29 | 0.63 | -6.81 | 2.92E-11 | 0.014 |
| M - NTC | Pseudobutyrivibrio | 2.55 | 0.92 | 2.78 | 2.82E-02 | 12.769 |
| M - NTC | Aerococcus | -3.37 | 0.62 | -5.47 | 1.80E-07 | 0.034 |
| M - NTC | Anaerosporobacter | -1.46 | 0.56 | -2.63 | 4.26E-02 | 0.231 |
| M - NTC | Victivallis | 4.10 | 0.83 | 4.95 | 2.80E-06 | 60.337 |
| M - ZnCu | Ruminococcaceae_unclassified | -0.41 | 0.14 | -2.94 | 1.73E-02 | 0.662 |
| M - ZnCu | Roseburia | -0.77 | 0.28 | -2.76 | 2.96E-02 | 0.464 |

| | | | | | | |
|---|---|---|---|---|---|---|
| M - ZnCu | Phascolarctobacterium | -0.76 | 0.28 | -2.75 | 3.06E-02 | 0.467 |
| M - ZnCu | Treponema | 2.59 | 0.67 | 3.90 | 5.49E-04 | 13.396 |
| M - ZnCu | Oscillibacter | -1.48 | 0.32 | -4.67 | 1.37E-05 | 0.229 |
| M - ZnCu | Porphyromonadaceae_unclassified | -1.10 | 0.32 | -3.43 | 3.69E-03 | 0.334 |
| M - ZnCu | Olsenella | 1.97 | 0.70 | 2.81 | 2.58E-02 | 7.198 |
| M - ZnCu | Butyricicoccus | 0.74 | 0.25 | 2.96 | 1.60E-02 | 2.094 |
| M - ZnCu | Ruminococcus | -0.52 | 0.20 | -2.68 | 3.68E-02 | 0.592 |
| M - ZnCu | Bifidobacterium | 2.14 | 0.71 | 3.02 | 1.35E-02 | 8.519 |
| M - ZnCu | Selenomonas | 2.64 | 0.65 | 4.06 | 3.01E-04 | 14.018 |
| M - ZnCu | Clostridia_unclassified | 1.95 | 0.45 | 4.37 | 6.93E-05 | 7.048 |
| M - ZnCu | Campylobacter | 2.37 | 0.72 | 3.31 | 5.13E-03 | 10.670 |
| M - ZnCu | Proteobacteria_unclassified | 1.44 | 0.55 | 2.61 | 4.50E-02 | 4.204 |
| M - ZnCu | Desulfovibrio | 3.48 | 0.85 | 4.07 | 2.94E-04 | 32.331 |
| M - ZnCu | Clostridium_XlVb | -1.53 | 0.44 | -3.50 | 2.63E-03 | 0.217 |
| M - ZnCu | Subdivision5_unclassified | 6.72 | 0.80 | 8.43 | 0.00E+00 | 825.986 |
| M - ZnCu | Anaerovibrio | -2.38 | 0.86 | -2.76 | 2.95E-02 | 0.093 |
| M - ZnCu | Fibrobacter | 4.24 | 0.91 | 4.66 | 1.29E-05 | 69.651 |
| M - ZnCu | Pseudoscardovia | 8.59 | 0.53 | 16.24 | 0.00E+00 | 5400.143 |
| M - ZnCu | Alphaproteobacteria_unclassified | 7.77 | 0.67 | 11.67 | 0.00E+00 | 2357.536 |
| M - ZnCu | Butyricimonas | 6.47 | 0.60 | 10.85 | 0.00E+00 | 647.544 |
| M - ZnCu | Acidaminococcaceae_unclassified | 2.46 | 0.91 | 2.71 | 3.40E-02 | 11.649 |
| M - ZnCu | Helicobacter | 4.04 | 0.69 | 5.82 | 1.94E-08 | 56.681 |
| M - ZnCu | Gammaproteobacteria_unclassified | 3.45 | 1.08 | 3.19 | 7.56E-03 | 31.366 |
| M - ZnCu | Sharpea | 7.36 | 0.73 | 10.03 | 0.00E+00 | 1569.047 |
| M - ZnCu | Bifidobacteriaceae_unclassified | 6.14 | 0.85 | 7.19 | 1.39E-12 | 463.919 |
| M - ZnCu | Oligosphaera | 6.38 | 0.61 | 10.43 | 0.00E+00 | 590.861 |
| M - ZnCu | Veillonella | 7.63 | 0.72 | 10.63 | 0.00E+00 | 2062.008 |
| M - ZnCu | Anaerotruncus | 3.86 | 1.04 | 3.71 | 1.11E-03 | 47.385 |
| M - ZnCu | Mucispirillum | 4.16 | 0.92 | 4.54 | 5.59E-05 | 63.958 |
| M - ZnCu | Betaproteobacteria_unclassified | 4.73 | 0.86 | 5.51 | 1.60E-07 | 113.178 |
| M - ZnCu | Staphylococcus | -4.44 | 0.77 | -5.75 | 8.59E-08 | 0.012 |
| M - ZnCu | Selenomonadales_unclassified | 2.91 | 0.93 | 3.14 | 9.28E-03 | 18.286 |
| M - ZnCu | Anaerobiospirillum | 5.48 | 1.03 | 5.33 | 3.25E-07 | 239.354 |
| M - ZnCu | Faecalitalea | 2.91 | 1.10 | 2.64 | 4.10E-02 | 18.286 |
| M - ZnCu | Anaerofustis | -4.90 | 0.83 | -5.92 | 1.10E-08 | 0.007 |
| M - ZnCu | Hydrogenoanaerobacterium | -4.43 | 1.12 | -3.97 | 4.30E-04 | 0.012 |
| M - ZnCu | Desulfovibrionales_unclassified | 4.35 | 0.35 | 12.44 | 0.00E+00 | 77.587 |
| M - ZnCu | Turicibacter | 3.15 | 0.83 | 3.78 | 9.22E-04 | 23.243 |
| M - ZnCu | Candidatus_Saccharibacteria_unclassified | 4.10 | 0.80 | 5.14 | 1.24E-06 | 60.265 |

| | | | | | | |
|---|---|---|---|---|---|---|
| M - ZnCu | Cellulosilyticum | -4.90 | 0.74 | -6.62 | 8.13E-11 | 0.007 |
| M - ZnCu | Actinomycetales_unclassified | 3.95 | 0.76 | 5.19 | 7.66E-07 | 51.817 |
| M - ZnCu | Romboutsia | -6.06 | 0.48 | -12.67 | 0.00E+00 | 0.002 |
| M - ZnCu | Pseudobutyrivibrio | 3.18 | 0.87 | 3.63 | 1.60E-03 | 23.927 |
| M - ZnCu | Victivallis | 4.10 | 0.79 | 5.20 | 1.13E-06 | 60.337 |
| M - ZnCu | Faecalicoccus | -2.74 | 0.67 | -4.07 | 2.64E-04 | 0.065 |
| M - ZnCu | Succiniclasticum | -6.25 | 0.74 | -8.41 | 0.00E+00 | 0.002 |
| M - ZnCu | Pyramidobacter | 5.15 | 0.75 | 6.83 | 1.90E-11 | 172.365 |
| M - ZnCu | Anaeroplasma | 4.51 | 0.95 | 4.74 | 1.23E-05 | 91.064 |
| NTC - AB | Streptococcus | 0.73 | 0.26 | 2.79 | 2.70E-02 | 2.077 |
| NTC - AB | Elusimicrobium | 7.41 | 1.20 | 6.16 | 3.36E-09 | 1647.648 |
| NTC - AB | Staphylococcus | 4.74 | 0.84 | 5.61 | 1.04E-07 | 114.191 |
| NTC - AB | Selenomonadales_unclassified | 4.45 | 1.01 | 4.39 | 5.61E-05 | 85.843 |
| NTC - AB | Flavonifractor | -5.70 | 1.06 | -5.40 | 1.73E-06 | 0.003 |
| NTC - AB | Anaerofustis | -4.43 | 0.91 | -4.89 | 3.68E-06 | 0.012 |
| NTC - AB | Candidatus_Saccharibacteria_unclassified | 4.88 | 0.87 | 5.59 | 8.49E-08 | 132.086 |
| NTC - AB | Butyrivibrio | -6.22 | 1.14 | -5.45 | 3.37E-07 | 0.002 |
| NTC - AB | Actinomycetales_unclassified | -3.83 | 0.83 | -4.60 | 1.91E-05 | 0.022 |
| NTC - AB | Spirochaetales_unclassified | 4.29 | 0.66 | 6.52 | 1.57E-10 | 72.866 |
| NTC - AB | Aerococcus | 3.37 | 0.64 | 5.24 | 5.81E-07 | 29.148 |
| NTC - AB | Victivallis | -3.67 | 0.86 | -4.24 | 1.26E-04 | 0.026 |
| NTC - AB | Bacillus | 4.29 | 1.17 | 3.65 | 1.46E-03 | 72.866 |
| NTC - AB | Anaeroplasma | 4.48 | 1.04 | 4.30 | 8.55E-05 | 88.324 |
| NTC - AB | Akkermansia | -5.70 | 0.66 | -8.57 | 0.00E+00 | 0.003 |
| NTC - ZnCu | Oscillibacter | -1.13 | 0.33 | -3.42 | 3.53E-03 | 0.323 |
| NTC - ZnCu | Porphyromonadaceae_unclassified | -1.22 | 0.34 | -3.65 | 1.45E-03 | 0.294 |
| NTC - ZnCu | Bifidobacterium | 2.53 | 0.74 | 3.40 | 3.92E-03 | 12.507 |
| NTC - ZnCu | Selenomonas | 2.44 | 0.68 | 3.57 | 1.91E-03 | 11.452 |
| NTC - ZnCu | Clostridia_unclassified | 1.73 | 0.47 | 3.68 | 1.27E-03 | 5.630 |
| NTC - ZnCu | Erysipelotrichaceae_unclassified | 1.73 | 0.58 | 3.00 | 1.44E-02 | 5.622 |
| NTC - ZnCu | Campylobacter | 2.30 | 0.75 | 3.06 | 1.17E-02 | 9.975 |
| NTC - ZnCu | Desulfovibrio | 4.48 | 0.90 | 5.00 | 5.72E-06 | 88.435 |
| NTC - ZnCu | Subdivision5_unclassified | 4.80 | 0.84 | 5.74 | 7.82E-08 | 120.998 |
| NTC - ZnCu | Fibrobacter | 5.10 | 0.96 | 5.33 | 4.06E-07 | 163.736 |
| NTC - ZnCu | Pseudoscardovia | 8.68 | 0.55 | 15.64 | 0.00E+00 | 5866.004 |
| NTC - ZnCu | Alphaproteobacteria_unclassified | 7.55 | 0.70 | 10.82 | 0.00E+00 | 1898.769 |
| NTC - ZnCu | Butyricimonas | 4.82 | 0.63 | 7.71 | 2.66E-14 | 123.961 |
| NTC - ZnCu | Acidaminococcaceae_unclassified | 2.47 | 0.95 | 2.60 | 4.59E-02 | 11.871 |
| NTC - ZnCu | Peptococcus | 0.98 | 0.32 | 3.05 | 1.25E-02 | 2.653 |

| | | | | | | |
|---|---|---|---|---|---|---|
| NTC - ZnCu | Helicobacter | 7.24 | 0.73 | 9.94 | 0.00E+00 | 1388.032 |
| NTC - ZnCu | Sharpea | 6.53 | 0.77 | 8.49 | 0.00E+00 | 686.438 |
| NTC - ZnCu | Bifidobacteriaceae_unclassified | 5.70 | 0.90 | 6.36 | 5.53E-10 | 298.908 |
| NTC - ZnCu | Oligosphaera | 5.28 | 0.64 | 8.22 | 6.66E-16 | 195.871 |
| NTC - ZnCu | Veillonella | 6.38 | 0.75 | 8.47 | 0.00E+00 | 589.486 |
| NTC - ZnCu | Mucispirillum | 5.87 | 0.96 | 6.11 | 4.35E-09 | 355.331 |
| NTC - ZnCu | Betaproteobacteria_unclassified | 5.15 | 0.90 | 5.73 | 4.30E-08 | 172.250 |
| NTC - ZnCu | Selenomonadales_unclassified | 4.45 | 0.97 | 4.58 | 2.49E-05 | 85.843 |
| NTC - ZnCu | Anaerobiospirillum | 4.48 | 1.08 | 4.15 | 1.71E-04 | 88.324 |
| NTC - ZnCu | Anaerococcus | -4.10 | 0.94 | -4.38 | 7.07E-05 | 0.017 |
| NTC - ZnCu | Faecalitalea | 4.95 | 1.15 | 4.29 | 1.31E-04 | 140.592 |
| NTC - ZnCu | Howardella | 2.56 | 0.96 | 2.66 | 3.89E-02 | 12.885 |
| NTC - ZnCu | Flavonifractor | -5.57 | 1.01 | -5.51 | 3.12E-07 | 0.004 |
| NTC - ZnCu | Anaerofustis | -4.90 | 0.87 | -5.65 | 4.82E-08 | 0.007 |
| NTC - ZnCu | Turicibacter | 2.46 | 0.87 | 2.82 | 2.47E-02 | 11.715 |
| NTC - ZnCu | Candidatus_Saccharibacteria_unclassified | 4.88 | 0.84 | 5.84 | 1.92E-08 | 132.086 |
| NTC - ZnCu | Butyrivibrio | -6.05 | 1.09 | -5.54 | 1.28E-07 | 0.002 |
| NTC - ZnCu | Cellulosilyticum | -4.90 | 0.78 | -6.31 | 2.06E-09 | 0.007 |
| NTC - ZnCu | Weissella | -4.72 | 1.08 | -4.36 | 9.94E-05 | 0.009 |
| NTC - ZnCu | Spirochaetales_unclassified | 4.29 | 0.63 | 6.81 | 5.62E-11 | 72.866 |
| NTC - ZnCu | Aerococcus | 3.37 | 0.62 | 5.47 | 1.47E-07 | 29.148 |
| NTC - ZnCu | Faecalicoccus | -2.21 | 0.71 | -3.13 | 9.52E-03 | 0.110 |
| NTC - ZnCu | Succiniclasticum | -6.25 | 0.78 | -8.01 | 2.33E-15 | 0.002 |
| NTC - ZnCu | Pyramidobacter | 5.64 | 0.79 | 7.13 | 2.17E-12 | 282.478 |
| NTC - ZnCu | Anaeroplasma | 4.48 | 1.00 | 4.49 | 3.14E-05 | 88.324 |

a) Delta Estimates were calculated using glht function in the multcomp package in R. The glht function calculates the estimates by subtracting the estimate of X2 (second treatment group in the column “Comparison”) from the estimate of X1 (first treatment group in the column “Comparison”). Estimates for each treatment group were obtained with Gamma glm models, which were used as the input for the glht function. See the provided R-script (electronic supplemental material) for the procedure.

b) Fold Changes were calculated by taking the exponential function from the Delta Estimate (due to the logarithmic link function that was used in with Gamma glm models). This was done in R using the command exp(Delta.Estimate). See the provided R-script (electronic supplemental material) for the procedure. Relative abundances of all genera are presented in Supplemental figure SX.

Table S3. Pairwise comparisons of negative binomial GLMs of abundances of each genera between treatment groups. Rarefied and subsampled OTU table was used as the input.

| Comparison ($X_1$ - $X_2$) | Genus | Delta Estimate (X1 - X2) [a] | Std. Error | z-value | p.adjusted | Fold Change (X1 / X2) [b] |
|---|---|---|---|---|---|---|
| AB - ZnCu | Veillonellaceae_unclassified | 1.21234387 | 0.33269944 | 3.64396126 | 0.00158131 | 3.36135402 |
| AB - ZnCu | Oscillibacter | -1.432626 | 0.35487338 | -4.0370062 | 0.00028535 | 0.23868132 |
| AB - ZnCu | Porphyromonadaceae_unclassified | -0.9640726 | 0.34852224 | -2.7661723 | 0.02893672 | 0.3813367 |
| AB - ZnCu | Streptococcus | -0.9078819 | 0.25697297 | -3.532986 | 0.00224082 | 0.40337772 |
| AB - ZnCu | Prevotellaceae_unclassified | 0.85321027 | 0.27317443 | 3.12331678 | 0.00965566 | 2.34716981 |
| AB - ZnCu | Selenomonas | 3.04217777 | 0.62717819 | 4.85057964 | 5.98E-06 | 20.9508197 |
| AB - ZnCu | Clostridia_unclassified | 1.58321472 | 0.45087436 | 3.5114321 | 0.00265948 | 4.87058824 |
| AB - ZnCu | Campylobacter | 2.1701959 | 0.63654643 | 3.40932851 | 0.00353628 | 8.76 |
| AB - ZnCu | Desulfovibrio | 3.07269331 | 0.87581187 | 3.50839423 | 0.0023786 | 21.6 |
| AB - ZnCu | Anaerovibrio | 0.44561218 | 0.10322547 | 4.31688194 | 8.89E-05 | 1.56144578 |
| AB - ZnCu | Peptococcus | 0.74193734 | 0.28442987 | 2.6085071 | 0.04490273 | 2.1 |
| AB - ZnCu | Oxalobacteraceae_unclassified | 1.53686722 | 0.44070683 | 3.48727797 | 0.00260496 | 4.65 |
| AB - ZnCu | Faecalicoccus | -2.3232044 | 0.61879187 | -3.7544197 | 0.00089624 | 0.09795918 |
| M - AB | Roseburia | -0.8614451 | 0.26223116 | -3.2850602 | 0.00542459 | 0.422551 |
| M - AB | Streptococcus | 1.04825084 | 0.25689019 | 4.08054053 | 0.00028992 | 2.852657 |
| M - AB | Prevotellaceae_unclassified | -1.1685243 | 0.27740881 | -4.2122825 | 0.0001068 | 0.31082529 |
| M - AB | Anaerovibrio | -2.7243866 | 0.25190262 | -10.815237 | 0 | 0.06558642 |
| M - AB | Pseudoramibacter | 2.89958841 | 1.10369578 | 2.62716273 | 0.04261924 | 18.1666667 |
| M - AB | Allisonella | 1.45990618 | 0.53051035 | 2.75189011 | 0.02926791 | 4.30555556 |
| M - AB | Oxalobacteraceae_unclassified | -1.3137237 | 0.41136664 | -3.1935591 | 0.00773409 | 0.2688172 |
| M - NTC | Prevotellaceae_unclassified | -0.8797698 | 0.27934219 | -3.1494341 | 0.00876736 | 0.4148784 |
| M - NTC | Anaerovibrio | -0.7830954 | 0.30180021 | -2.5947478 | 0.04116923 | 0.45698925 |
| M - ZnCu | Ruminococcaceae_unclassified | -0.4012112 | 0.13492098 | -2.9736755 | 0.01527879 | 0.66950864 |
| M - ZnCu | Faecalibacterium | -0.7493485 | 0.26744944 | -2.8018323 | 0.02602334 | 0.47267442 |
| M - ZnCu | Roseburia | -0.7794952 | 0.25026243 | -3.1147113 | 0.01029925 | 0.45863747 |
| M - ZnCu | Treponema | 2.79662947 | 0.74336141 | 3.76213968 | 0.00107451 | 16.389313 |
| M - ZnCu | Oscillibacter | -1.4855684 | 0.3382575 | -4.3918271 | 7.11E-05 | 0.22637363 |
| M - ZnCu | Porphyromonadaceae_unclassified | -1.1365224 | 0.33263108 | -3.4167654 | 0.00337963 | 0.32093317 |
| M - ZnCu | Olsenella | 1.98922401 | 0.56687383 | 3.50911246 | 0.00252894 | 7.30985915 |
| M - ZnCu | Butyricicoccus | 0.76490792 | 0.25902375 | 2.95304164 | 0.0165773 | 2.1487965 |
| M - ZnCu | Ruminococcus | -0.5667333 | 0.16511636 | -3.4323265 | 0.00327959 | 0.56737589 |
| M - ZnCu | Bifidobacterium | 2.2042704 | 0.79761532 | 2.7635758 | 0.02915979 | 9.06363636 |
| M - ZnCu | Selenomonas | 2.66563313 | 0.59947487 | 4.4466136 | 7.13E-05 | 14.3770492 |
| M - ZnCu | Clostridia_unclassified | 1.9029851 | 0.43098873 | 4.41539411 | 0.00021892 | 6.70588235 |
| M - ZnCu | Campylobacter | 2.45958884 | 0.60834529 | 4.04308029 | 0.00039292 | 11.7 |
| M - ZnCu | Desulfovibrio | 2.60268969 | 0.86865068 | 2.99624435 | 0.01392845 | 13.5 |
| M - ZnCu | Clostridium_XlVb | -1.540445 | 0.50212119 | -3.067875 | 0.01084537 | 0.21428571 |

| | | | | | | |
|---|---|---|---|---|---|---|
| M - ZnCu | Clostridium_XIVa | -1.6094379 | 0.60403501 | -2.6644779 | 0.03826044 | 0.2 |
| M - ZnCu | Anaerovibrio | -2.2787744 | 0.25465513 | -8.9484725 | 0 | 0.10240964 |
| M - ZnCu | Parabacteroides | -1.3581235 | 0.47611974 | -2.8524831 | 0.0219854 | 0.25714286 |
| M - ZnCu | Gammaproteobacteria_unclassified | 4.78749174 | 1.65951315 | 2.88487726 | 0.02014325 | 120 |
| M - ZnCu | Faecalicoccus | -2.793208 | 0.67528503 | -4.1363393 | 0.00015585 | 0.06122449 |
| NTC - AB | Streptococcus | 0.74675663 | 0.26842882 | 2.78195402 | 0.02801263 | 2.11014493 |
| NTC - AB | Anaerovibrio | -1.9412912 | 0.19206673 | -10.107379 | 0 | 0.14351852 |
| NTC - ZnCu | Oscillibacter | -1.0977336 | 0.35265113 | -3.1128031 | 0.00987795 | 0.33362637 |
| NTC - ZnCu | Porphyromonadaceae_unclassified | -1.262393 | 0.34951026 | -3.6118912 | 0.00155403 | 0.28297604 |
| NTC - ZnCu | Olsenella | 1.8412201 | 0.59382959 | 3.10058665 | 0.01049971 | 6.30422535 |
| NTC - ZnCu | Bifidobacterium | 2.59172065 | 0.83583121 | 3.10077036 | 0.01024605 | 13.3527273 |
| NTC - ZnCu | Selenomonas | 2.49469466 | 0.62772358 | 3.97419299 | 0.00037095 | 12.1180328 |
| NTC - ZnCu | Clostridia_unclassified | 1.82644503 | 0.44739085 | 4.08243713 | 0.00023809 | 6.21176471 |
| NTC - ZnCu | Erysipelotrichaceae_unclassified | 1.82866403 | 0.54885298 | 3.33179208 | 0.00498575 | 6.22556391 |
| NTC - ZnCu | Campylobacter | 2.35327821 | 0.63594609 | 3.70043661 | 0.00122548 | 10.52 |
| NTC - ZnCu | Desulfovibrio | 3.29583687 | 0.87263445 | 3.77688146 | 0.00076604 | 27 |
| NTC - ZnCu | Anaerovibrio | -1.495679 | 0.1956628 | -7.6441664 | 4.92E-14 | 0.22409639 |
| NTC - ZnCu | Peptococcus | 1.08856195 | 0.2769759 | 3.9301685 | 0.00054426 | 2.97 |
| NTC - ZnCu | Faecalicoccus | -1.9177393 | 0.54732992 | -3.5038086 | 0.00241794 | 0.14693878 |

a) Delta Estimates were calculated using glht function in the multcomp package in R. The glht function calculates the estimates by subtracting the estimate of X2 (second treatment group in the column “Comparison”) from the estimate of X1 (first treatment group in the column “Comparison”). Estimates for each treatment group were obtained with Negative binomial glm models, which were used as the input for the glht function. See the provided R-script (electronic supplemental material) for the procedure.

b) Fold Changes were calculated by taking the exponential function from the Delta Estimate (due to the logarithmic link function that was used in with negative binomial glm models). This was done in R using the command exp(Delta.Estimate). See the provided R-script (electronic supplemental material) for the procedure. Relative abundances of all genera are presented in Supplemental figure SX.

Table S4. Pairwise comparisons of gamma distribution GLMs of relative abundances of each ARG or MGE between treatment groups.

| Comparison ($X_1$ - $X_2$) | Gene | Delta Estimate (X1 - X2) [a] | Std. Error | z-value | p.adjusted | Fold Change (X1 / X2) [b] |
|---|---|---|---|---|---|---|
| AB – ZnCu | tetM | -1.9609786 | 0.56562766 | -3.4669071 | 0.00294252 | 0.14072065 |
| AB - ZnCu | blaSFO | -1.6624443 | 0.6172798 | -2.6931779 | 0.03566898 | 0.18967479 |
| AB - ZnCu | tet(32) | -0.8667544 | 0.29147371 | -2.9736966 | 0.01532269 | 0.42031351 |
| AB - ZnCu | tetA | -3.3801931 | 0.80640588 | -4.1916772 | 0.00012203 | 0.03404088 |
| AB - ZnCu | aac(6)-Iy | -2.2241923 | 0.64701255 | -3.4376339 | 0.00316533 | 0.10815474 |
| AB - ZnCu | orf37-IS26 | 1.70692972 | 0.27849818 | 6.12905162 | 2.66E-09 | 5.51201204 |
| AB - ZnCu | aac3ia | 1.64899726 | 0.62668685 | 2.63129385 | 0.04215185 | 5.20176117 |
| AB - ZnCu | aadA17 | 3.03054434 | 0.75440944 | 4.01710817 | 0.00036428 | 20.7085019 |
| AB - ZnCu | aph(3)-ia | 1.6305567 | 0.38277192 | 4.25986499 | 0.00014771 | 5.10671682 |
| M - AB | erm(A) | -2.1113086 | 0.76595471 | -2.7564406 | 0.0297697 | 0.12107941 |
| M - AB | dfrA15 | 2.48130513 | 0.70164466 | 3.53641277 | 0.00223079 | 11.9568595 |
| M - AB | ISEfm1-Entero | -2.9992904 | 0.98225854 | -3.0534633 | 0.01213616 | 0.04982241 |
| M - AB | mcr-1 | 4.2850184 | 0.79531544 | 5.38782244 | 2.35E-07 | 72.6038824 |
| M - AB | IS1247 | -2.8561563 | 0.93354463 | -3.0594749 | 0.01153335 | 0.0574893 |
| M - AB | tet(32) | 0.9720145 | 0.3057002 | 3.17963315 | 0.00818034 | 2.64326396 |
| M - AB | vanRB | 5.93955828 | 0.87592439 | 6.78090298 | 2.52E-11 | 379.767142 |
| M - AB | bacA | -1.8909589 | 0.62089051 | -3.0455593 | 0.01254414 | 0.15092702 |
| M - AB | cmr | -2.9181191 | 0.81540135 | -3.5787519 | 0.00198127 | 0.05403523 |
| M - AB | IS1111 | 1.07880591 | 0.4100313 | 2.63103307 | 0.04210773 | 2.94116543 |
| M - AB | tetW | 0.59294255 | 0.11487403 | 5.16167619 | 1.79E-06 | 1.80930456 |
| M - AB | cmlA1 | 2.9568897 | 0.5889674 | 5.02046413 | 2.50E-06 | 19.2380425 |
| M - AB | ermT | 2.28000567 | 0.54303264 | 4.19865312 | 0.00013153 | 9.77673588 |
| M - AB | tetA | 4.40984486 | 0.84576562 | 5.2140271 | 1.32E-06 | 82.2567008 |
| M - AB | orf37-IS26 | -1.7069297 | 0.29209135 | -5.8438214 | 2.38E-08 | 0.18142196 |
| M - AB | cmlA5 | 3.00497288 | 0.59048343 | 5.08900462 | 2.59E-06 | 20.1856687 |
| M - AB | QnrB4 | 1.78597148 | 0.32353984 | 5.52009763 | 1.68E-07 | 5.96537237 |
| M - AB | mphA | -2.1385063 | 0.53931174 | -3.9652507 | 0.00035192 | 0.11783072 |
| M - AB | aac(3)-Xa | 3.47371474 | 0.36875121 | 9.42021236 | 0 | 32.256344 |
| M - AB | ant6-ia | -0.8613593 | 0.21763422 | -3.9578302 | 0.00049996 | 0.42258728 |
| M - AB | Aac6-Aph2 | 1.5327412 | 0.30740967 | 4.98598883 | 4.25E-06 | 4.63085353 |
| M - AB | aph(3)-ia | -1.6305567 | 0.40145458 | -4.0616219 | 0.00025999 | 0.19582053 |
| M - AB | tetC | -1.9095796 | 0.69594238 | -2.743876 | 0.03098174 | 0.14814265 |
| M - AB | acc3-iva | -2.5851379 | 0.72497665 | -3.5658223 | 0.00199241 | 0.07538568 |
| M - NTC | dfra21 | -3.9631864 | 0.7464272 | -5.3095418 | 4.30E-07 | 0.01900247 |
| M - NTC | mcr-1 | 3.11136771 | 0.79531544 | 3.9121178 | 0.00054159 | 22.4517308 |
| M - NTC | IncF | -2.7468676 | 0.71209149 | -3.8574644 | 0.00068427 | 0.06412843 |
| M - NTC | IS613 | -0.5945431 | 0.19546268 | -3.0417219 | 0.01234019 | 0.55181463 |

| | | | | | | |
|---|---|---|---|---|---|---|
| M - NTC | int1-a-marko | -4.0443799 | 0.74755476 | -5.4101454 | 3.42E-07 | 0.01752057 |
| M - NTC | vanYB | -1.6261959 | 0.40225007 | -4.0427485 | 0.00037319 | 0.19667634 |
| M - NTC | IS1247 | -2.5124873 | 0.93354463 | -2.6913414 | 0.03556148 | 0.08106635 |
| M - NTC | blaSFO | -2.8016459 | 0.64740851 | -4.3274777 | 0.00012122 | 0.06071006 |
| M - NTC | vanHB | -1.8205826 | 0.46912614 | -3.8807956 | 0.00056227 | 0.16193138 |
| M - NTC | vanRB | 4.19033987 | 0.87592439 | 4.78390594 | 1.58E-05 | 66.0452339 |
| M - NTC | IS200 | -2.1107617 | 0.78217107 | -2.6985934 | 0.03530454 | 0.12114566 |
| M - NTC | sugE | -2.2099317 | 0.67741053 | -3.2623227 | 0.00604713 | 0.10970814 |
| M - NTC | IS1111 | 1.07880591 | 0.4100313 | 2.63103307 | 0.04215758 | 2.94116543 |
| M - NTC | dfrA12 | -3.5203626 | 0.73837227 | -4.7677341 | 9.18E-06 | 0.0295887 |
| M - NTC | tetW | 0.31333733 | 0.11487403 | 2.72766025 | 0.03253207 | 1.36798291 |
| M - NTC | cmlA1 | 2.9568897 | 0.5889674 | 5.02046413 | 2.29E-06 | 19.2380425 |
| M - NTC | aac3ia | 1.89166903 | 0.65727471 | 2.8780493 | 0.02077131 | 6.63042586 |
| M - NTC | cmlA5 | 3.00497288 | 0.59048343 | 5.08900462 | 3.01E-06 | 20.1856687 |
| M - NTC | QnrB4 | 1.78597148 | 0.32353984 | 5.52009763 | 9.52E-08 | 5.96537237 |
| M - NTC | mphA | -1.4728425 | 0.53931174 | -2.7309668 | 0.03232183 | 0.22927286 |
| M - NTC | fabK | 5.66404178 | 1.09149277 | 5.18926182 | 1.53E-06 | 288.311582 |
| M - NTC | aac(3)-Xa | 3.47371474 | 0.36875121 | 9.42021236 | 0 | 32.256344 |
| M - NTC | ant6-ia | -0.7219647 | 0.21763422 | -3.3173308 | 0.0050286 | 0.48579688 |
| M - NTC | Aac6-Aph2 | 1.5327412 | 0.30740967 | 4.98598883 | 3.46E-06 | 4.63085353 |
| M - NTC | aadA17 | 4.68465257 | 0.7912313 | 5.92071191 | 9.04E-09 | 108.272649 |
| M - NTC | strA | 3.30889729 | 0.98127942 | 3.37202353 | 0.00437277 | 27.3549443 |
| M - NTC | spcN | -1.9089525 | 0.49374891 | -3.8662415 | 0.0006272 | 0.14823558 |
| M - NTC | acc3-iva | -2.1937907 | 0.72497665 | -3.0260156 | 0.01318488 | 0.11149331 |
| M - ZnCu | dfrA15 | 2.48130513 | 0.70164466 | 3.53641277 | 0.00217154 | 11.9568595 |
| M - ZnCu | ISEfm1-Entero | -3.2048101 | 0.98225854 | -3.262695 | 0.00593547 | 0.04056661 |
| M - ZnCu | mdth | 2.9246563 | 0.86237088 | 3.39141356 | 0.00380529 | 18.6278225 |
| M - ZnCu | erm(B) | 0.93109457 | 0.33870954 | 2.74894695 | 0.03070325 | 2.53728489 |
| M - ZnCu | mcr-1 | 4.2850184 | 0.79531544 | 5.38782244 | 3.78E-07 | 72.6038824 |
| M - ZnCu | IS613 | -0.5225922 | 0.19546268 | -2.6736165 | 0.03762232 | 0.5929814 |
| M - ZnCu | vat(E) | -1.2091678 | 0.37578186 | -3.2177387 | 0.00677412 | 0.29844553 |
| M - ZnCu | tetM | -1.743174 | 0.59323529 | -2.9384193 | 0.01727917 | 0.17496418 |
| M - ZnCu | blaSFO | -1.6624443 | 0.64740851 | -2.5678444 | 0.04998973 | 0.18967479 |
| M - ZnCu | tetU | 2.29801572 | 0.69439174 | 3.30939381 | 0.00522197 | 9.95441055 |
| M - ZnCu | vanRB | 5.93955828 | 0.87592439 | 6.78090298 | 4.37E-11 | 379.767142 |
| M - ZnCu | bacA | -2.1036734 | 0.62089051 | -3.3881552 | 0.00401755 | 0.12200742 |
| M - ZnCu | cmr | -2.1813192 | 0.81540135 | -2.6751478 | 0.03746989 | 0.11289251 |
| M - ZnCu | IS1111 | 1.07880591 | 0.4100313 | 2.63103307 | 0.04184856 | 2.94116543 |
| M - ZnCu | tetW | 0.39580496 | 0.11487403 | 3.44555648 | 0.00324454 | 1.48557954 |

| | | | | | | |
|---|---|---|---|---|---|---|
| M - ZnCu | cmlA1 | 2.9568897 | 0.5889674 | 5.02046413 | 3.46E-06 | 19.2380425 |
| M - ZnCu | ermT | 2.12523835 | 0.54303264 | 3.91364754 | 0.000564 | 8.37489337 |
| M - ZnCu | aac(6)-Iy | -2.2241923 | 0.67859249 | -3.2776553 | 0.00590438 | 0.10815474 |
| M - ZnCu | aac3ia | 1.89166903 | 0.65727471 | 2.8780493 | 0.02082748 | 6.63042586 |
| M - ZnCu | cmlA5 | 3.00497288 | 0.59048343 | 5.08900462 | 3.37E-06 | 20.1856687 |
| M - ZnCu | QnrB4 | 1.78597148 | 0.32353984 | 5.52009763 | 1.33E-07 | 5.96537237 |
| M - ZnCu | mphA | -2.0535919 | 0.53931174 | -3.8078013 | 0.00084003 | 0.12827332 |
| M - ZnCu | aac(3)-Xa | 3.47371474 | 0.36875121 | 9.42021236 | 0 | 32.256344 |
| M - ZnCu | ant6-ia | -0.8695084 | 0.21763422 | -3.9952742 | 0.00036814 | 0.41915756 |
| M - ZnCu | Aac6-Aph2 | 1.5327412 | 0.30740967 | 4.98598883 | 4.42E-06 | 4.63085353 |
| M - ZnCu | aadA17 | 4.68465257 | 0.7912313 | 5.92071191 | 8.40E-09 | 108.272649 |
| M - ZnCu | tetC | -1.958219 | 0.69594238 | -2.8137659 | 0.02511551 | 0.14110952 |
| NTC - AB | ere(A) | -3.7985065 | 1.118659 | -3.3955892 | 0.00364477 | 0.02240421 |
| NTC - AB | dfra21 | 3.96318637 | 0.71169041 | 5.56869437 | 5.08E-07 | 52.6247409 |
| NTC - AB | erm(A) | -3.4796663 | 0.73030916 | -4.7646483 | 1.12E-05 | 0.03081769 |
| NTC - AB | IncF | 2.74686756 | 0.6789526 | 4.04574274 | 0.00033225 | 15.5937089 |
| NTC - AB | int1-a-marko | 4.04437991 | 0.7127655 | 5.67420832 | 5.89E-08 | 57.0757829 |
| NTC - AB | vanYB | 1.62619587 | 0.3835304 | 4.24007036 | 0.00011458 | 5.0844958 |
| NTC - AB | vat(E) | -0.9633283 | 0.35829394 | -2.6886537 | 0.03557084 | 0.38162061 |
| NTC - AB | blaSFO | 2.80164588 | 0.6172798 | 4.53869688 | 3.12E-05 | 16.471735 |
| NTC - AB | vanHB | 1.82058264 | 0.44729422 | 4.07021273 | 0.0002616 | 6.17545545 |
| NTC - AB | VanB | 1.4812986 | 0.56054483 | 2.64260507 | 0.04109909 | 4.39865408 |
| NTC - AB | bacA | -1.8909589 | 0.59199588 | -3.1942096 | 0.00729479 | 0.15092702 |
| NTC - AB | sugE | 2.20993173 | 0.6458856 | 3.42155287 | 0.00316531 | 9.11509408 |
| NTC - AB | cmr | -2.9181191 | 0.77745468 | -3.7534266 | 0.00096566 | 0.05403523 |
| NTC - AB | dfrA12 | 3.52036262 | 0.70401033 | 5.0004417 | 2.61E-06 | 33.7966817 |
| NTC - AB | ermT | 1.43379945 | 0.5177613 | 2.76922867 | 0.02844954 | 4.19460614 |
| NTC - AB | tetA | 2.23750725 | 0.80640588 | 2.77466635 | 0.02830619 | 9.36994519 |
| NTC - AB | orf37-IS26 | -1.7069297 | 0.27849818 | -6.1290516 | 2.37E-09 | 0.18142196 |
| NTC - AB | aac3ia | -1.6489973 | 0.62668685 | -2.6312938 | 0.04216712 | 0.19224258 |
| NTC - AB | aadD | -2.5570934 | 0.82041991 | -3.1168105 | 0.01000558 | 0.07752976 |
| NTC - AB | fabK | -6.4623226 | 1.04069753 | -6.209607 | 1.87E-08 | 0.00156117 |
| NTC - AB | aadA17 | -3.0305443 | 0.75440944 | -4.0171082 | 0.00035953 | 0.04828935 |
| NTC - AB | aph(3)-ia | -1.6305567 | 0.38277192 | -4.259865 | 0.00014469 | 0.19582053 |
| NTC - AB | spcN | 1.9089525 | 0.47077111 | 4.05494826 | 0.00026519 | 6.74601867 |
| NTC - ZnCu | dfra21 | 3.96318637 | 0.71169041 | 5.56869437 | 1.47E-07 | 52.6247409 |
| NTC - ZnCu | erm(A) | -2.2898479 | 0.73030916 | -3.1354501 | 0.00921967 | 0.10128186 |
| NTC - ZnCu | mdth | 3.19193363 | 0.82223837 | 3.8820052 | 0.00065477 | 24.3354377 |
| NTC - ZnCu | erm(B) | 0.84582312 | 0.32294688 | 2.61907817 | 0.04367295 | 2.32989481 |

| | | | | | | |
|---|---|---|---|---|---|---|
| NTC - ZnCu | IncF | 2.74686756 | 0.6789526 | 4.04574274 | 0.00030468 | 15.5937089 |
| NTC - ZnCu | int1-a-marko | 4.04437991 | 0.7127655 | 5.67420832 | 1.30E-07 | 57.0757829 |
| NTC - ZnCu | vanYB | 1.62619587 | 0.3835304 | 4.24007036 | 0.00011774 | 5.0844958 |
| NTC - ZnCu | vat(E) | -1.2481406 | 0.35829394 | -3.4835659 | 0.00276248 | 0.28703803 |
| NTC - ZnCu | vanHB | 1.82058264 | 0.44729422 | 4.07021273 | 0.00022799 | 6.17545545 |
| NTC - ZnCu | VanB | 1.4812986 | 0.56054483 | 2.64260507 | 0.04103194 | 4.39865408 |
| NTC - ZnCu | tetU | 2.15358493 | 0.66207655 | 3.25277333 | 0.00610526 | 8.61568974 |
| NTC - ZnCu | IS200 | 2.11076166 | 0.74577086 | 2.83030859 | 0.02401747 | 8.25452605 |
| NTC - ZnCu | bacA | -2.1036734 | 0.59199588 | -3.5535271 | 0.00222767 | 0.12200742 |
| NTC - ZnCu | sugE | 2.20993173 | 0.6458856 | 3.42155287 | 0.00365646 | 9.11509408 |
| NTC - ZnCu | cmr | -2.1813192 | 0.77745468 | -2.8057187 | 0.02591365 | 0.11289251 |
| NTC - ZnCu | dfrA12 | 3.52036262 | 0.70401033 | 5.0004417 | 3.54E-06 | 33.7966817 |
| NTC - ZnCu | aadA2 | 1.89169834 | 0.72517442 | 2.60861151 | 0.04472002 | 6.6306202 |
| NTC - ZnCu | aac(6)-Iy | -2.2241923 | 0.64701255 | -3.4376339 | 0.00315928 | 0.10815474 |
| NTC - ZnCu | aadD | -2.6220232 | 0.82041991 | -3.1959527 | 0.00703458 | 0.07265572 |
| NTC - ZnCu | fabK | -5.6767979 | 1.04069753 | -5.454801 | 2.60E-07 | 0.00342451 |
| NTC - ZnCu | spcN | 1.9089525 | 0.47077111 | 4.05494826 | 0.00026009 | 6.74601867 |
| NTC - ZnCu | aph(2)-Id | -0.4948545 | 0.19239134 | -2.5721245 | 0.04936651 | 0.60965962 |

a) Delta Estimates were calculated using glht function in the multcomp package in R. The glht function calculates the estimates by subtracting the estimate of X2 (second treatment group in the column “Comparison”) from the estimate of X1 (first treatment group in the column “Comparison”). Estimates for each treatment group were obtained with Gamma glm models, which were used as the input for the glht function. See the provided R-script (electronic supplemental material) for the procedure.

b) Fold Changes were calculated by taking the exponential function from the Delta Estimate (due to the logarithmic link function that was used in with Gamma glm models). This was done in R using the command exp(Delta.Estimate). See the provided R-script (electronic supplemental material) for the procedure. Relative abundances of all genera are presented in Supplemental figure SX.

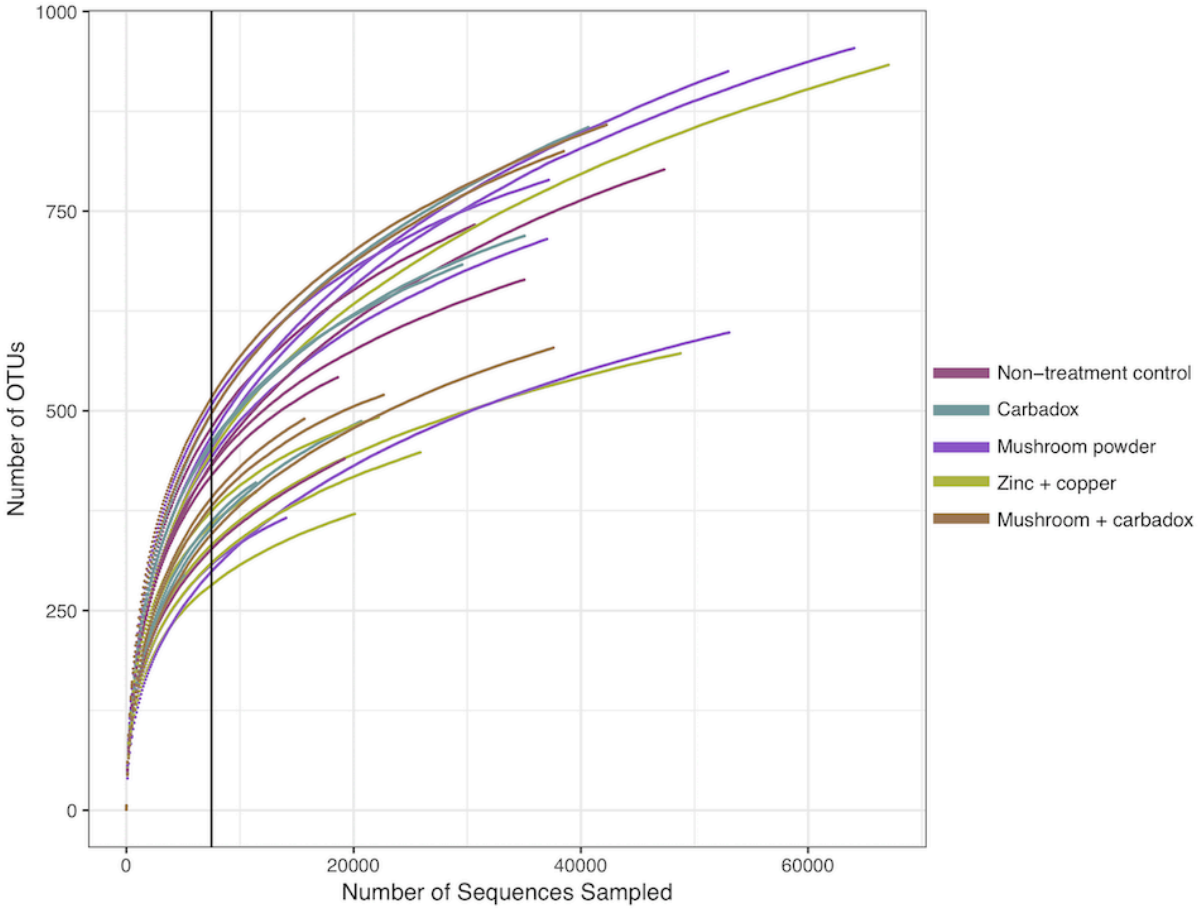


Fig. S4. Rarefaction curves. OTU collection curves determined from sequence analysis. Each line represents one sample. Vertical line shows the subsampling cutoff: 7500 sequences

Table S5. Assays that had unspecific amplification. Ct values in the negative control and mean Ct-values in samples.

| Assay | Ct in the negative control | Mean Ct in samples |
|---|---|---|
| 16S old 1_1 | 24.14393 | 11.10064 |
| blaOXY-1_1118 | 23.06278 | 22.71912 |
| cmlV_911 | 26.30517 | 25.85088 |
| czcA_1536 | 16.75728 | 17.97052 |
| fabK_1520 | 25.9036 | 19.75017 |
| intI1F165_clinical_359 | 25.72579 | 23.82255 |
| tetPA_1507 | 24.76249 | NA |
